# Quantification and demonstration of the constriction-by-rachet mechanism in the dynamin molecular motor

**DOI:** 10.1101/2020.09.10.289546

**Authors:** Oleg Ganichkin, Renee Vancraenenbroeck, Gabriel Rosenblum, Hagen Hofmann, Alexander S. Mikhailov, Oliver Daumke, Jeffrey K. Noel

## Abstract

Dynamin oligomerizes into helical filaments on tubular membrane templates and, through constriction, cleaves them in a GTPase-driven way. Structural observations of GTP-dependent cross-bridges between neighboring filament turns have led to the suggestion that dynamin operates as a molecular ratchet motor. However, the proof of such mechanism remains absent. Particularly, it is not known whether a powerful enough stroke is produced and how the motor modules would cooperate in the constriction process. Here, we characterized the dynamin motor modules by single molecule (sm) FRET and found strong nucleotide-dependent conformational changes. Integrating smFRET with molecular dynamics simulations allowed us to determine the forces generated in a power stroke. Subsequently, the quantitative force data and the measured kinetics of the GTPase cycle were incorporated into a model including both a dynamin filament, with explicit motor cross-bridges, and a realistic deformable membrane template. In our simulations, collective constriction of the membrane by dynamin motor modules, based on the ratchet mechanism, is directly reproduced and analyzed. Functional parallels between the dynamin system and actomyosin in the muscle are seen. Through concerted action of the motors, tight membrane constriction to the hemifission radius can be reached. Our experimental and computational study provides an example of how collective motor action in megadalton molecular assemblies can be approached and explicitly resolved.

## 1 Introduction

Dynamin is a mechano-chemical GTPase that plays a central role in clathrin-mediated endocytosis (CME) [1–4]. The 100kD protein polymerizes into helical filaments that coil around the necks of vesicles budding from the membrane. In presence of GTP, such filaments constrict and cut the necks, thus allowing vesicles to separate. This process is crucial for cellular nutrient uptake and for synaptic transmission. Mutations in dynamin are associated with severe neurodegenerative disease and muscular disorders [5]. Structural studies have detailed the dynamin filament structure on membrane tubes and observed the filament’s ability to form nucleotide-dependent cross-bridges between its neighboring turns [6, 7]. These studies, in tandem with biophysical experiments showing torque generation by dynamin filaments in the presence of GTP, have led to proposals that dynamin functions as a molecular ratchet motor [8–10]. Confirming this hypothesis requires a molecular-level understanding of the principal GTP-dependent motor function and showing that the candidate powerstroke [6] can provide sufficient force to drive membrane constriction. So far, theoretical studies of dynamin concentrated on the elastic properties of dynamin filaments [11–13] and on the effects exhibited by passive elastic filaments on the membranes [14, 15]. The non-equilibrium motor activity of dynamin, based on GTP hydrolysis, was only treated in a phenomenological way [16, 17].

Dynamin consists of five domains (Figure 1A) [18, 19]. Its stalks polymerize into a helical filament whose elementary units are criss-cross stalk dimers. The filament is anchored to the membrane by pleckstrin homology (PH) domains connected by flexible linkers to the stalk. GTPase (G) domains are connected to the filament via bundle signaling elements (BSE). The disordered C-terminal proline-rich domain (PRD) is involved in recruitment to membrane necks. Hinge 1 and hinge 2 form flexible joints between BSE and the stalk and between the G domain and BSE, respectively. We refer to the combination of the G domain and BSE as the *motor module* (MM) of dynamin (Figure 1A) [6, 20, 21].

**Figure 1.**
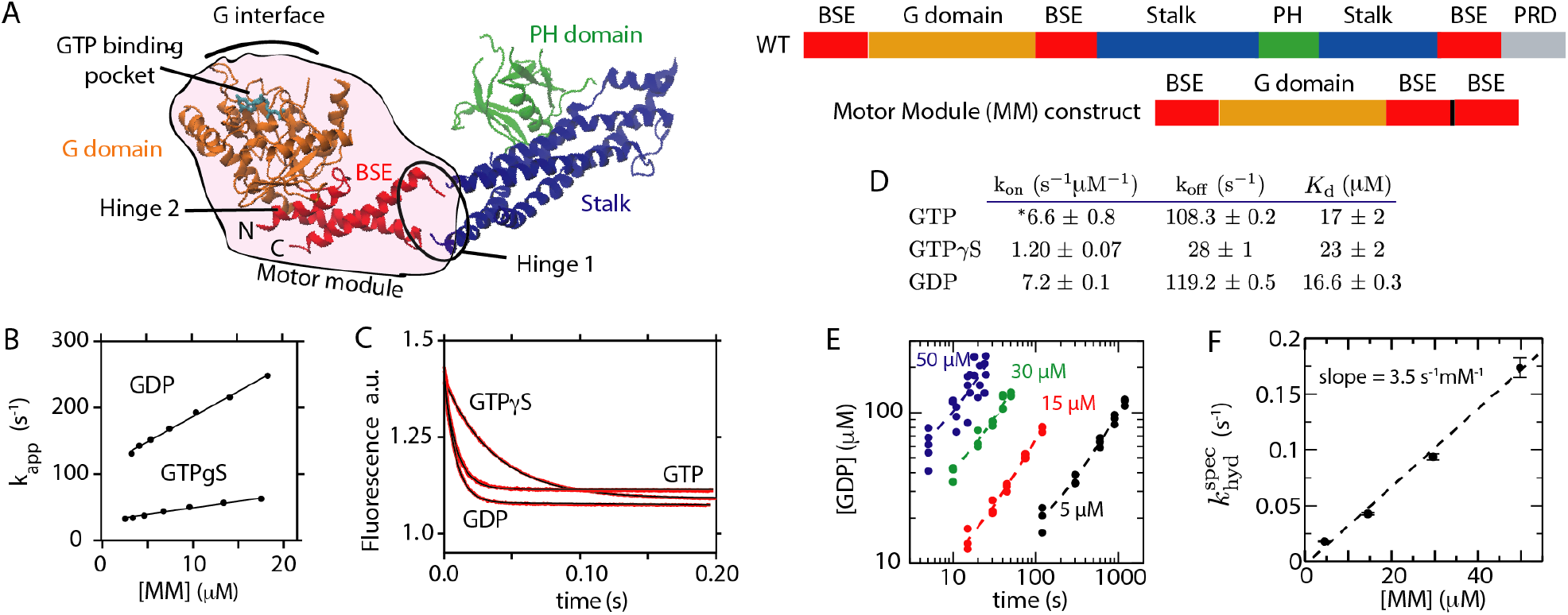
Kinetic characterization of the dynamin motor module. A) The dynamin monomer has four structurally characterized domains (PDB 3SNH [18]). The motor module (MM) contains the G domain and the BSE. A continuous MM construct was created by fusing the third helix of the BSE onto the N-terminal portion of the dynamin sequence. B) Association rate constants are obtained from the slope of *k*_app_ versus MM concentration. See Figure S1 for mant-GTP association curves. C) Mant-nucleotide dissociation rates were measured in stopped-flow experiments by mixing with excess unlabeled nucleotide. D) Table of determined mant-nucleotide binding/dissociation rates of the MM construct. K_d_ is defined by the ratio k_off_ */*k_on_. *\**GTP on rate value is determined from kinetic modeling of GTPase activity (see Section S1.1.2) because mant binding did not show a single exponential fluorescence increase for mant-GTP (Figure S1). E) GDP production as a function of time in the linear regime for increasing concentrations of MM and an initial GTP concentration of 1 mM. Dotted lines are linear fits to three independent experiments. F) Slope from (E) divided by the protein concentration gives the specific hydrolysis rate 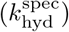. Linear dependence of the specific hydrolysis rate on protein concentration indicates that GTPase activity is controlled by dimerization.

When dynamin is assembled into a helical filament, G domains in adjacent rungs are optimally oriented for cross-dimerization [7], which can explain the enhancement of GTPase activity in the presence of membranes [21]. Moreover, the conformation of dynamin’s MM is sensitive to its nucleotide state. In the presence of the non-hydrolyzable GTP analog *β,γ*-methyleneguanosine 5^*1*^-triphosphate (GMPPCP), the MM crystallized in an open BSE conformation [6], whereas a closed conformation of the BSE was observed in the nucleotide-free state or when bound to GDP or to 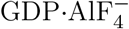, a mimic of the transition state of GTP hydrolysis [18,19,21,22]. While this structural shift is the obvious candidate for the power stroke driving motor function, it remains unknown how strong it is, since even small energy preferences might result in the different structural states captured in the available structures. For example, a dynamin family member, MxA, shows open and closed states in crystallography [23], but only weak preferences in solution [24].

In this study, kinetic measurements for the GTPase cycle of the dynamin MM are first performed. After that, we explore nucleotide-dependent conformational changes in the MM using single-molecule FRET experiments. Their results, in conjunction with molecular modeling, are then used to quantify the forces generated by single dynamin motors and to refine the ratchet effect. Next, we incorporate the determined forces and the measured kinetics into a polymer-like computational model that resolves individual dynamin motors and includes a deformable membrane template with lipid flows within it. Direct simulations of the model reproduce collective tight constriction of membrane necks down to the hemi-fission radius by dynamin filaments in the presence of GTP.

## 2 Results

In order to study the kinetic and energetic parameters of a single dynamin motor, we have worked with a MM construct containing only the G domain and BSE [21] (Figure 1A). This allows clean interpretation of nucleotide on and off rates, as it removes the complicating process of tetramers (or higher oligomers) associating via their G interfaces. Additionally, using the MM construct for the smFRET experiments allows us to ensure we are capturing the intrinsic G domain/BSE conformational ensemble.

### 2.1 Exploring dynamin’s GTPase cycle

Nucleotide binding and dissociation rates to this construct were evaluated using stopped-flow experiments with fluorescently-labeled mant-nucleotides (Figures 1B,C and S1A). The affinities for GTP and GDP were both in the low micromolar range (Figure 1D). Therefore, at physiological nucleotide concentrations of GTP (300 µM) and GDP (30 µM) [25], about 90% of monomeric MMs in the cell are bound to GTP. Previous measurements [26, 27] reporting even higher GTP affinity were performed using full-length dynamin, thus also including possible contributions from nucleotide-induced assembly that enhance nucleotide occupancy.

A low basal rate of GTPase hydrolysis and a linear dependence of the specific hydrolysis rate 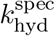 on MM concentration were observed (Figure 1E,F and Section S1.1.1), confirming that MM dimerization should precede the hydrolysis and is required for it. These GTPase measurements additionally indicated that the GTP-bound dimer has a weak affinity 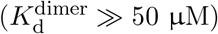.

Running the GTPase assay beyond the linear regime allows GDP to compete with GTP for the active site. Therefore, the GTPase kinetics of such an experiment contain information on the relative affinity between GTP and GDP for the MM. A kinetic model was devised to fit experimental traces and determine unknown parameters (SI Section S1.1.2). Varying this setup with different initial conditions (Figures S1B,D) allowed us to estimate an apparent GTP on-rate of 6.6 s^−1^µM ^−1^, which was not straightforward to determine with mant-GTP (Figure S1A).

Measuring nucleotide exchange kinetics within the dimer is complicated by the low MM dimerization affinity. To resolve this difficulty, we engineered a Zn^2+^-dependent metal bridge by introducing histidines into the G interface (Figure S2A). In line with our predictions, addition of Zn^2+^ to this MM-HH construct induced dimerization in gel-filtration (Fig. S2B) and FRET experiments (Fig. S2C). Of note, addition of GTP*γ*S reduced dimer formation (Fig. S2C). X-ray crystallography confirmed that Zn^2+^ stabilized a similar dimer arrangement as previously observed in the GDP-bound MM dimer [22] (Figure S2A and Table S2).

Dimer dissociation rates were measured by FRET experiments and stopped-flow. Addition of GDP slowed the dimer dissociation rate of the Zn^2+^-stabilized MM-HH dimer six-fold relative to the apo state (Figure S2D,E) consistent with a long-lived post-hydrolysis dimeric state. Addition of GTP greatly accelerated dimer dissociation, suggesting that the GTP-bound state is not compatible with the engineered Zn^2+^ bridge. Relevant to the motor cycle, mant-GDP did not bind to empty Zn^2+^-stabilized MM-HH dimers. Furthermore, mant-GDP dissociation from MM-HH dimers was 100-fold slower in the presence of Zn^2+^ (Figure S2G-J). These observations imply that nucleotide exchange is greatly reduced in the MM dimer, therefore simplifying dynamin’s motor cycle (see below).

### 2.2 Nucleotide-dependent motor module conformation in solution

To characterize nucleotide-dependent conformational changes in the MM, single-molecule (sm) FRET was employed. We monitored the relative orientation of the G domain and BSE by introducing a pair of FRET dyes, one in BSE and the other in the G domain (Figures 2A,B and S3). The experiments were performed on freely diffusing MMs.

**Figure 2.**
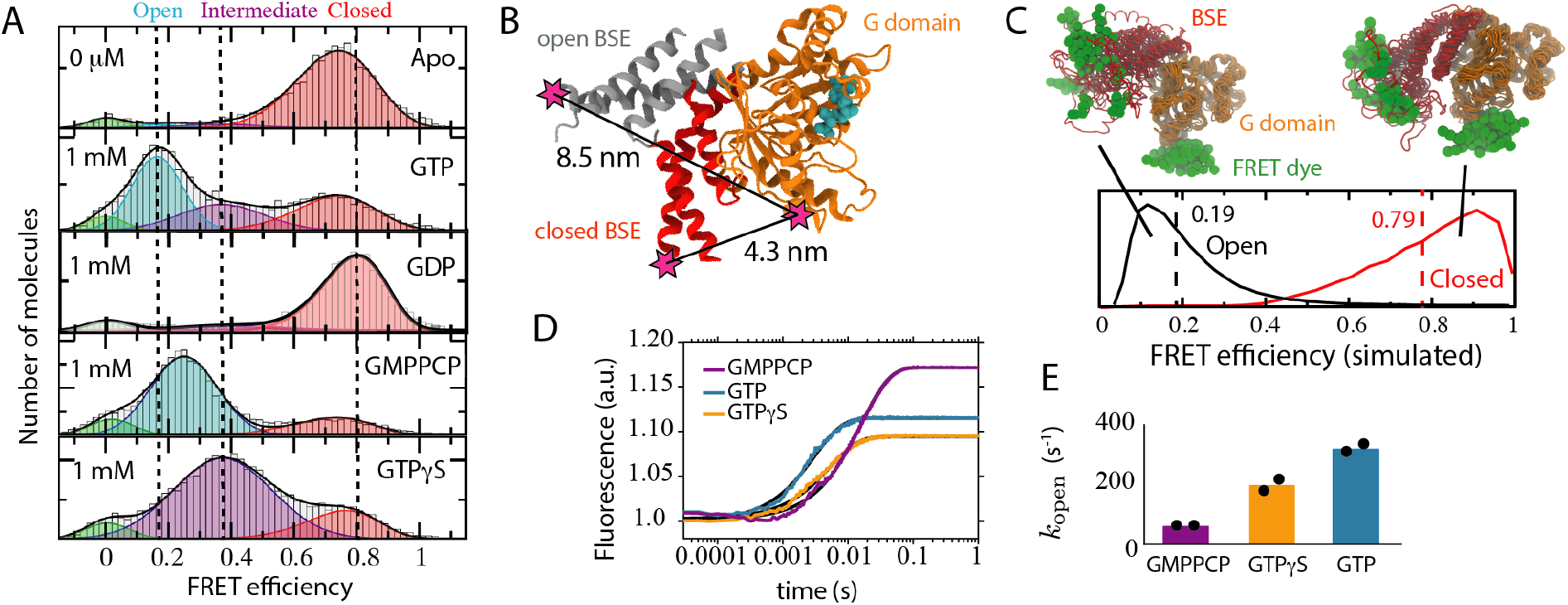
FRET measurements reveal the dependence of the G domain/BSE conformation on nucleotide state. A) smFRET histograms were fit to a four Gaussian state model and were colored according to the approximate position of the peaks: red, closed; purple, intermediate open; cyan, open; green, molecules lacking an active acceptor dye. Apo and GDP states peak at high FRET while the other ligands show heterogeneity. The picomolar concentrations of MM used for smFRET ensured that the ensembles corresponded to MM monomers. Distance between FRET labeling sites based on the crystallographic open (GMPPCP-bound [6]) and closed (GDP-bound [22]) BSE orientations. C) Dotted lines (solid lines) show the average FRET (FRET distribution) from molecular simulations based on the open or closed crystal structures. The close correspondence of the average FRET with the smFRET data provide evidence that the crystal structures describe the high and low FRET states seen in smFRET. Several snapshots from the two simulation ensembles are displayed with their G domains aligned and FRET dye represented as green spheres. D) Kinetics of opening were measured by mixing apo, doubly-labeled MM with saturating concentrations (1 mM) of nucleotide. Single exponential fits to donor fluorescence are shown. E) The opening rate determined by averaging the single exponential rate constants from two independent experiments (shown as black dots). The measured rate involves two processes: nucleotide binding and the conformational change.

In the presence of GTP, three populations could be discerned (Figures 2A and S3C) by fitting to Gaussian distributions: a dominant open state with a FRET efficiency peak (*E*) at *E* = 0.18, an intermediate state at *E* = 0.39, and a closed state at *E* = 0.77. The peak values of the open and closed states were in excellent agreement with the predicted mean FRET efficiencies from simulations based on the crystal structures of 0.19 and 0.79, respectively (Figure 2C). The GTP analogs (GMPPCP and GTP*γ*S) showed bimodal distributions. Interestingly, the low FRET peak for GTP*γ*S coincided with the intermediate peak found in the presence of GTP. Note that the non-hydrolyzable GTP analogues produce different structural ensembles from those found with GTP. Care has therefore to be taken when interpreting experiments using GTP analogs. Remarkably, in absence of nucleotides or in presence of GDP, the conformational distributions were strongly shifted towards the closed state. For the apo protein, the position of the FRET peak coincided with that of the GTP-bound closed state. For GDP, the closed state was shifted slightly higher to *E* = 0.8, consistent with crystal data (Figure S3D).

Ensemble stopped-flow measurements were used to obtain relaxation rates to the equilibrium distributions. The rate of opening, i.e. of a transition from apo (closed) to the GTP-bound state, was fitted to a single exponential with a time constant of approximately 300 s^−1^ (Figure 2D,E). This rate contains both nucleotide binding and the conformational change. However, since nucleotide binding at 1 mM GTP is much faster than opening, the measured rate must correspond to the conformational change. Because 300 s^−1^ is large as compared to the overall turnover rate of the GTPase cycle, GTP binding and opening are combined in our summary of the GTPase cycle (Figure 3). The rate of closing in solution, induced by GTP dissociation, is larger than 105 s^−1^ (Figure S3E). Although, this parameter is of little relevance for dynamin’s function since, in the filament, closing takes place within a dimer and under load.

**Figure 3.**
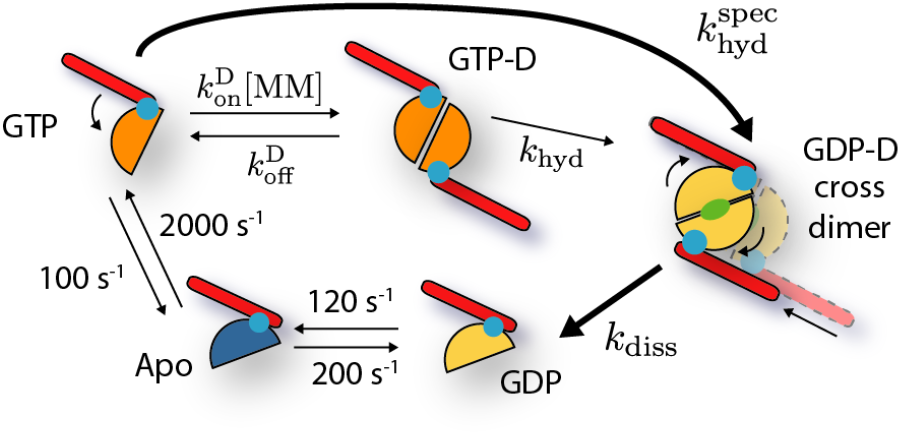
GTPase cycle for the MM. Ligand binding kinetics are taken from Figure 1 assuming physiological concentrations of 300 µM GTP and 30 µM GDP. [MM] refers to the concentration of GTP-bound MM. Experimental measurements of GTPase activity in the presence of membranes set a lower limit of 4 s^−1^ [18] for the whole cycle.

### 2.3 Operation cycle of the dynamin motor

The kinetic and conformational results are summarized in the diagram of the GTPase cycle (Figure 3). The GTP-bound MM monomers are predominantly open and must dimerize (GTP-D) to enable hydrolysis. Hydrolysis leads to a strong preference in the MM dimer (GDP-D) for the closed state. The MM dimer in its GDP·Pi or GDP-bound states dissociates only slowly, consistent with the high stability of the dimers bound to 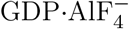 [21] and with our kinetic results with GDP (Figure S2E). Nucleotide exchange within the MM dimer is slow and can therefore be neglected in the scheme (Figure S2I,J). Once monomeric, GDP exchanges in favor of GTP, thus completing the cycle.

Note that hydrolysis (*k*_hyd_) and post-hydrolysis MM dimer dissociation (*k*_diss_) rates, as well as the dimerization 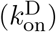 and dissociation 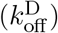 rates of MM monomers in the GTP-bound state, remain yet unknown, meaning the entire GTPase cycle could not be precisely specified. Therefore, in our model simulations, various values of the effective hydrolysis rate 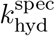 and of *k*_diss_ were chosen and explored (see Section S1.3 for details). Measurements of GTPase activity in the presence of membranes set a lower limit of 4 s^−1^ [18] for its turnover rate. The slowest measured rate in the forward direction is that of GDP dissociation at 120 s^−1^, yielding an upper limit for the overall turnover rate.

### 2.4 Determination of generated forces

In the helical filament, a pair of GTP-bound MMs belonging to adjacent rungs can dimerize to form a *cross-bridge* (Figure 4A). Upon hydrolysis in both MMs, the equilibrium conformation of the dimer shifts from the open to the closed ensemble thus shortening its equilibrium bridging length and creating a contracting force along it.

**Figure 4.**
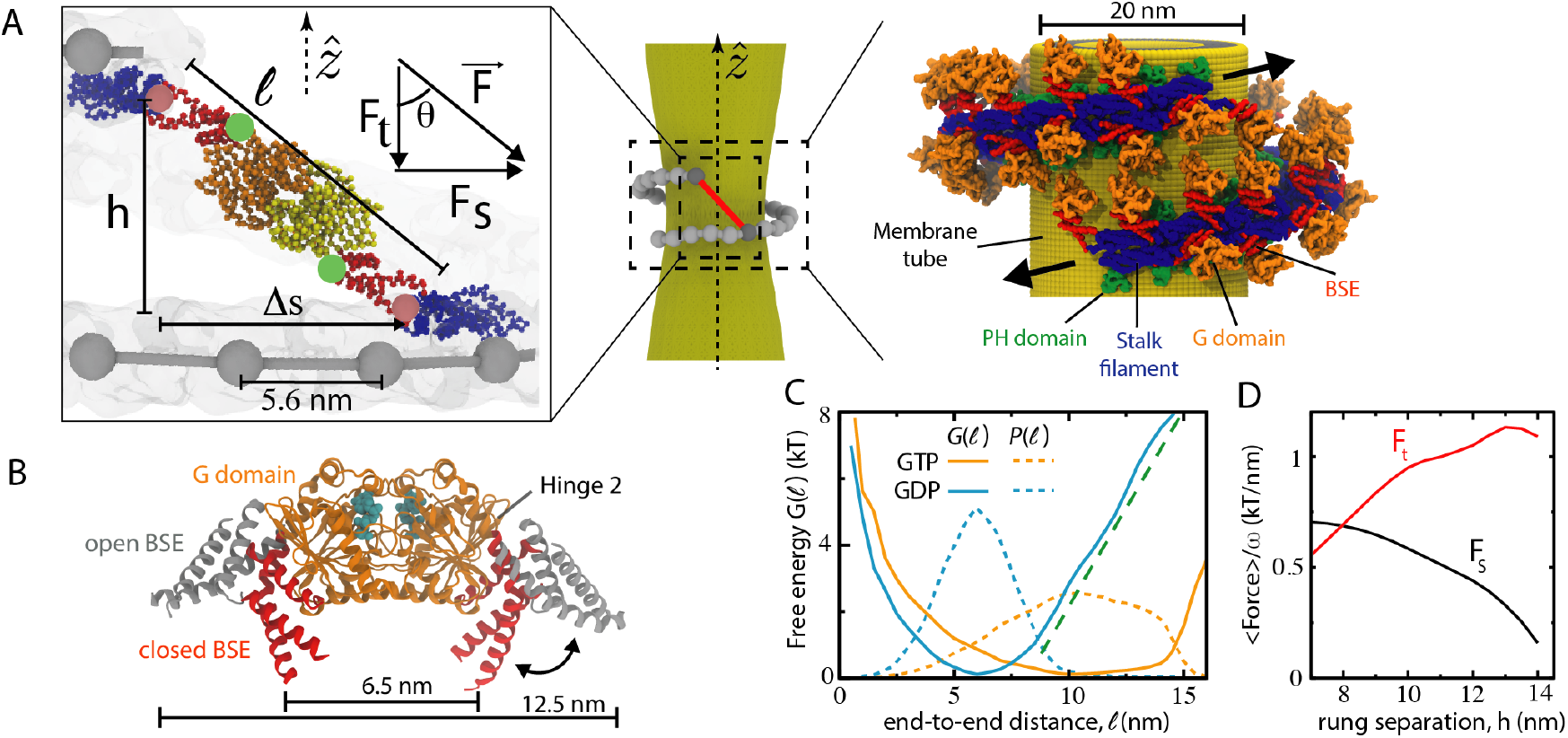
Determination of forces. A) (Center) An MM dimer forms a cross-bridge (red) between two neighboring filament rungs. (Left) Definition of cross-bridge geometry. The length *𝓁* is defined as the distance between hinge 1 connections (pink circles), green circles indicate hinges 2. (Right) A cartoon representation based on the cryo-EM structure of the “non-constricted” filament [28] shows the overall architecture of the dynamin filament. B) Overlay of open and closed G domain dimer crystal structures. Estimates for *𝓁*, i.e. the distance between hinge 1 attachment points, are shown. C) Free energies *G*_GTP_(*𝓁*) (orange) and *G*_GDP_(*𝓁*) (blue) obtained by MD simulations for MM dimers, the respective distributions *P*_GTP_(*𝓁*) and *P*_GDP_(*𝓁*) of distances are shown by orange and blue dashed lines. The dotted line has a slope of *F*_0_ = 1.3 k_B_T/nm (5.2 pN). D) Mean stall forces along the filament (black) and in the transverse direction (red) as functions of the separation *h* between the rungs (see SI for details).

The length of a bridging MM dimer is quantified by the end-to-end length distributions *P*_GTP_(*𝓁*) and *P*_GDP_(*𝓁*). Although distance distributions can be yielded by smFRET, our FRET experiments are limited to MM monomers. Therefore, probability distributions over *𝓁* for the two nucleotide states were obtained from MD simulations of a dual-basin structure-based model [29] (see Figure S5 and Section S1.2 for a detailed discussion). The MM dimer representation was built by connecting two MM monomers, each described by the dual-basin model, at their G interface (Figure 4B). Importantly, the simulations incorporated the experimentally-observed energetics by enforcing that the simulated FRET distribution in each MM monomer match the observed smFRET distribution of open and closed states (Figure S5).

The free energy profile of a GTP-bound MM dimer along *𝓁* shows a single broad basin (Figure 4C). This implies that initial cross-bridges can form over a wide range of distances. This property is enabled by the heterogeneity of the GTP-bound conformations (Figure 2A). In contrast, the free energy *G*_GDP_ for GDP-bound MM dimers has a single narrow minimum at a shorter *𝓁* (Figure 4C). The effective contracting force along *𝓁* is given by *F* (*𝓁*) = −*k*_B_*T* (*dG*_GDP_(*𝓁*)*/d𝓁*). In the simulations of membrane constriction, this instantaneous force, produced by a MM dimer when it is in the GDP-bound state and has length *𝓁*, is employed. Remarkably, the force remains approximately constant (see Figure 4C) at *F*_0_ = 1.3 k_B_T/nm over the range of *𝓁* from 8 nm to 14 nm.

11

In order to transform the cyclic changes of the GTPase cycle (Figure 3) into a directed force acting on the helical filament, a ratchet mechanism must be employed. The principal feature of the ratchet is a differential response to the power stroke and the recovery stroke. The transition from an open to a closed conformation, representing a power stroke, takes place when the two MMs from the neighboring rungs are dimerized and forming a cross-bridge. The recovery stroke, corresponding to the reverse transition to the open state, however, only occurs in the monomeric MM state, i.e. when the bridge is absent. Therefore, force is applied to the filament only in one part of the GTPase cycle.

There is however an additional component to the ratchet effect. If it were equally probable that the cross-bridges slanted to the right and to the left, shortening of such links could not have generated a net force. Hence, a directional bias in slanting is needed. For dynamin’s right-handed helix, preferentially slanting the upper MM to the right and the lower MM to the left with respect to the stalk filament would generate a constricting torque (i.e. as in Figure 4A). Indeed, the comparison of all available dynamin structures has shown that hinge 1 is biased in this way, and our MD simulations have indicated that the biased orientation is robust to thermal fluctuations (Figure S4).

One can calculate the mean force generated by a pair of MMs over many GTPase cycles by averaging the force applied between two immobilized stalk filaments (under stall conditions). Note that time averaging is equivalent to statistical averaging over an ensemble. Using the distributions for *𝓁* from MD simulations and the geometry of Figure 4C, the component of the mean force acting along the filament is 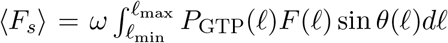 where *θ* is the slant angle and *ω* is the probability (often referred to as the *duty ratio* [30]) to find a MM in the dimerized state. Because of the ratchet property of hinge 1, only configurations with positive displacements Δ*s* are included into the average. The transverse component ⟨*F*_*t*_⟩ is given by the same equation with sin *θ* replaced by − cos *θ*.

The mean forces ⟨*F*_*s*_⟩ and ⟨*F*_*t*_⟩ are shown as functions of the rung separation *h* in Figure 4D. Note that the derived estimates are sensitive to the relative weights of the open and closed conformations in a GDP-bound MM (Figure S5C,D). Since the observed frequency of the open conformation in smFRET is likely enhanced by experimental noise, they should be viewed as providing the lower bounds for the force.

The torque, locally applied to the filament by a MM dimer motor, is ⟨*F*_*s*_⟩*R* where *R* is the filament radius. Assuming that a MM is found predominantly in the dimerized state, i.e. *ω* is near unity, (*F*_*s*_) is 2.9 pN (0.7 k_B_T/nm) and the mean torque generated by a single motor in the non-constricted state (with *h* = 8 nm and *R* = 25 nm as in Figure 4A) is 72 pN·nm.

### 2.5 Collective operation of motor modules and constriction of membranes

The MM dimers, connecting neighboring rungs, produce forces that tend to rotate them with respect to one another. Collectively, they generate the torque applied to the filament. We directly demonstrate this via computer simulations of a mesoscopic model. It was obtained by augmenting the recently developed coarse-grained description of a dynamin polymer filament on a deformable membrane tube [13] to additionally account for the GTPase motor activity of dynamin. In the model, MM dimers are introduced as elastic cross-bridges that connect the polymer beads, and apply forces between them. Since each bead corresponds to a dynamin stalk dimer, it has two MMs associated with it. The nucleotide state of each MM is tracked and the GTPase cycle is implemented within it. Dimerization, i.e. formation of a cross-bridge, can occur if both MMs are GTP-bound. After the hydrolysis-induced conformational change, the cross-bridge becomes strained and produces, as outlined above, a contracting force *F* (*𝓁*) for a cross-bridge of instantaneous length *𝓁*. The MM dimers dissociate (and thus cross-bridges disappear) at rate *k*_diss_. See Methods and Figure S6 for the details.

In endocytosis, constriction and cleavage of membrane necks is performed by short dynamin filaments with only a few tens of MMs [31]. Filaments with 28 or 40 beads were therefore probed in our simulations. They were carried out for various MM dimer dissociation rates *k*_diss_ (Video S1), thus yielding the regimes with different duty ratios *ω* (Eq. S8). Figure 5A gives example snapshots of the initial and final states. The dependence of the final steady-state ILR on the duty ratio is displayed in Figure 5B. Tight constriction from 10.5 nm to 2 − 3 nm is found above *ω* = 0.65. Increasing the filament length from 28 to 40 significantly decreases the ILR, however increasing the length beyond 40 beads does not lead to further decreases in ILR.

**Figure 5.**
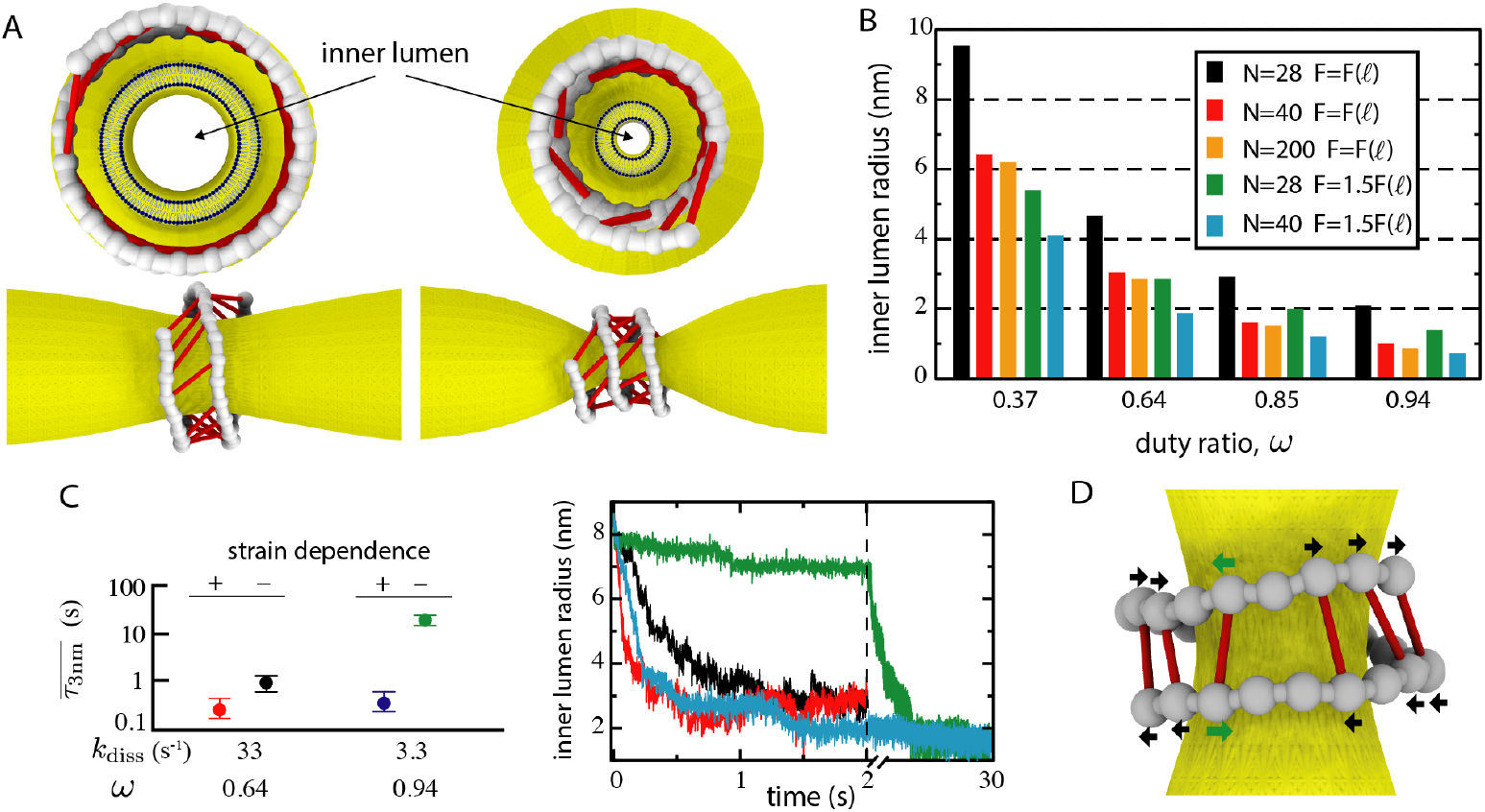
Membrane constriction by the dynamin oligomer. A) Snapshots of the initial (left) and final steady states for a filament with *N* = 40 dimerxs and duty ratio *ω* = 0.75. Cross-bridges indicated as red links. The tube is constricted from ILR of 10.5 nm to 2.5 nm. B) Dependence of the steady-state ILR on the duty ratio *ω* for different filament lengths *N* and motor strengths *F* (*𝓁*). Since the force depends on the population ratio of open and closed FRET states in the GDP-bound ensemble which can be affected by experimental noise, it may well be larger than in Figure 4C. The membrane has stiffness of 24 k_B_T (6×10^−20^ J) and tension of 0.03 k_B_T/nm^2^. All simulations start from thermal equilibrium in absence of GTP. C) Example of a configuration including a blocking link (with green arrows). Bridging GDP-bound MM dimers are shown as red links. D) Constriction is much accelerated under a strain-dependent MM dimer dissociation rate (green and black: strain-independent *k*_diss_, blue and red: strain-dependent *k*_diss_). (left) Average times 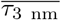 to reach an ILR of 3 nm; horizontal bars indicate the upper and lower quartiles over 50 constriction simulations. (right) Examples of time dependence of the ILR (the same color code as to the left).

Imaging of single endocytic pits during CME suggests that dynamin is active from a few seconds to tens of seconds before scission occurs [31]. Consistent with this, simulations showed that ILR below 3 nm could be reached within about a second for the duty ratios of 0.6 − 0.7 and within 15 s for *ω* = 0.94 (Figure 5C). Examination revealed that constriction speed was limited by *k*_diss_. When MM dimer dissociation is slow (corresponding to large duty ratios), cross-bridges continue to stay after the power stroke and are driven along with the sliding filament to reverse their slants (Figure 5D). The mis-oriented cross-bridges get oppositely stretched and block the constriction.

In the actomyosin system, key kinetic steps are sensitive to strain [32–35] and similar sensitivity may be present in dynamin. In our model, strain control could be implemented by introducing immediate dissociation of MM dimers once their slant is reversed and they become stretched. For a 40-bead filament and *ω* = 0.94, this reduced the time of constriction to 3 nm from 15 s down to 300 ms (Figure 5C and Video S2). Note that strain control only affects the kinetics, since the standing filament in the steady state cannot drive cross-bridges to alter their slants (and therefore the same ILR dependence as in Figure 5B holds). This effect could underlie the reported fast endocytosis within hundreds of milliseconds [36].

In *in vitro* experiments, scission of membrane tubes by long dynamin filaments can be observed. Surprisingly, such experiments find that breaking of the tube takes place not in the middle of the dynamin coat, but rather at its flanks [37, 38]. In line with this finding, our simulations show that constriction by a long filament begins near its ends (Figure 6A,B and Video S3) and only later propagates toward the middle. Fission may thus occur at a flank before the steady state has been reached. A simple explanation is that, in the middle of a long filament, a rung is pulled in opposite directions by its two neighboring rungs (Figures 6C,D). Such cancellation is absent for the terminal turns and, thus, only the motors on the flanks of a long helical filament apply a net torque and contribute to constriction of the membrane. Hence, the work produced by a long dynamin filament becomes independent of its length. Note that, in the employed model, dynamin-coated membrane tubes are assumed to be straight. Therefore, the effects of supercoiling, reported for very long tubes [17, 39], could not be considered here.

**Figure 6.**
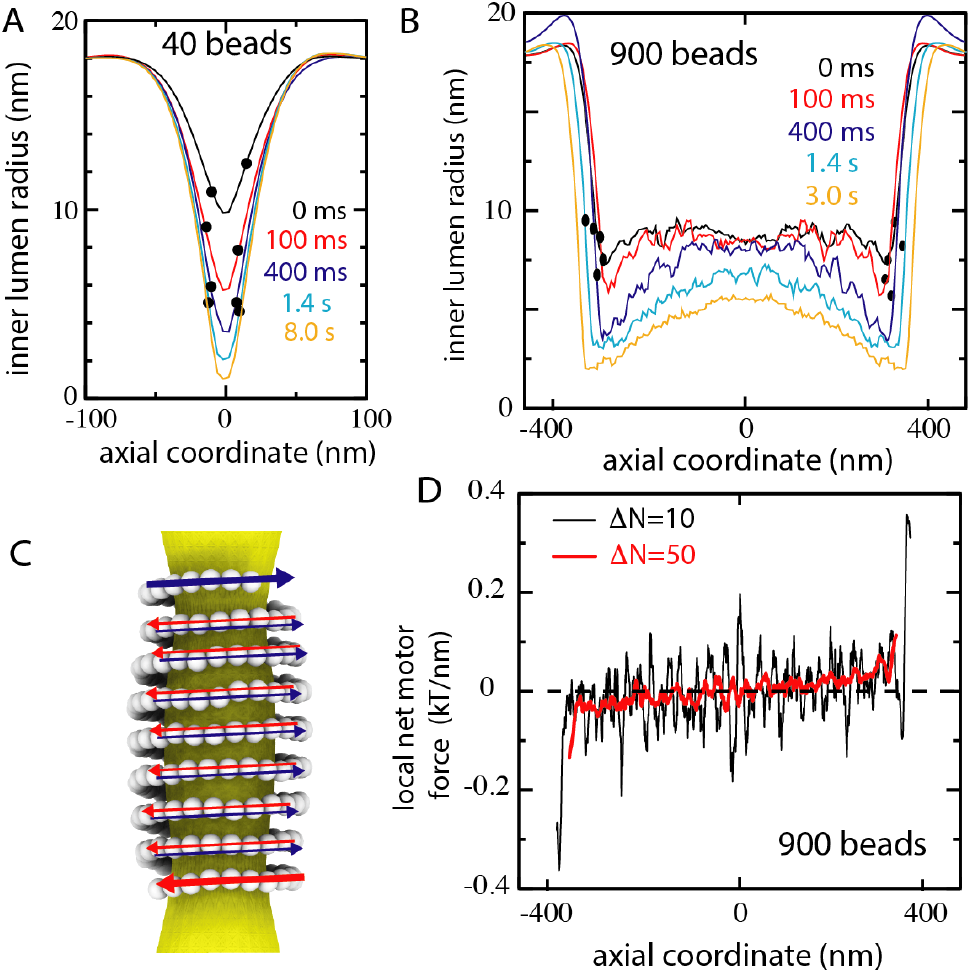
Long filaments first constrict near their ends. A,B) Evolutions of ILR profile under strain control for short (*N* = 40) and long (*N* = 900) filaments. Circles mark the positions of terminal beads. The membrane has stiffness of 24 k_B_T (6×10^−20^ J), tension of 0.03 k_B_T/nm^2^, and *ω* = 0.94. C) Motor forces in a dynamin coat tend to cancel each another, except for the flanks. Figure shows an *N* = 200 filament and cross-bridges are not shown. D) Local net motor force acting on the long filament along its direction. Running averages over Δ*N* = 10 (black) and Δ*N* = 50 (red) beads, averaged additionally over the first 500 ms, from the simulation in B are displayed.

## 3 Discussion

Since its discovery in the early nineties [40,41], dynamin has become the paradigm for studying GTPase-driven membrane remodeling. Biophysical experiments and structural studies have revealed dynamin to be a helical-shaped, torque-producing motor. In the current study, we provide the quantitative basis for the proposed ratchet operation and describe in detail the molecular mechanism whereby dynamin’s motor modules collectively induce filament sliding. Our smFRET experiments showed strong nucleotide-dependent shifts, which, in conjunction with molecular modeling, allowed the us to estimate the force generated by a single cross-bridge. Then, by incorporating these interactions into an energetically-realistic filament-on-a-membrane model, we directly demonstrate that through cooperative action, dynamin’s cross-bridges generate sufficient torque to drive strong constriction of biomembrane tubes. Importantly, the principal experimental features of membrane constriction by dynamin are either reproduced in our study or consistent with its results. Dynamin’s ability to create torque over a dynamin helix [17] is shown to arise from repeated cycles of interaction between neighboring MMs. It is confirmed that short dynamin helices with 1.5 to 2 turns are sufficient to produce tight constriction [31]. The fission at the edge of a dynamin coat, observed in the experiments with fluorescent membrane tubes pulled from giant unilamellar vesicles [37] or for nanotubes imaged by high-speed atomic force microscopy [38] could be explained by us. Typical durations of dynamin scission events [31, 36] are in agreement with the theoretic predictions.

The force generated by a single cross-bridge is much less than the overall force dynamin has been experimentally measured to produce [37] and less than the force that is theoretically required for strongly constricting the membrane tube (Figure S6E). Therefore, strong constriction requires that, at any time, most of the possible cross-bridges be formed (i.e. a high duty ratio) in order to cooperatively generate a large torque. Note that this cooperation does not come from synchronizing GTPase activity between MMs. Importantly, our simulations have shown that relative sliding of neighboring turns is possible even when MMs are working asynchronously and with large duty ratios. Sliding can be expedited with a strain-dependent dissociation rate.

We noted that nucleotide-dependent conformational changes are much more pronounced in dynamin than in the related MxA protein [24]. Whereas for MxA, the ratio of populations of the open to closed states in the presence of GDP was 0.43 (i.e., 30% to 70%), the GDP-bound open state was practically undetectable for dynamin (i.e. a ratio *<* 0.05). Thus, the resulting forces generated are at least three times stronger for dynamin. This highlights the functional difference of dynamin with MxA: MxA works as an antiviral machine that most likely acts on viral nucleocapsids rather than in the remodeling of membranes [42]. We therefore expect that the functions of dynamin superfamily members are modulated by tuning the nucleotide-dependent conformational preferences of the MM.

In our simulations, the filament is treated as an elastic ribbon whose local orientation with respect to the membrane remains fixed, even though it has been noticed that tilt deformations changing the orientation might contribute to filament breakup [43]. It is also assumed that nucleotide-driven conformational changes are limited to the MM. Moreover, axial symmetry of the membrane is assumed. The membrane is treated in the elastic Helfrich approximation, lipid flows within it are conserved. The filament sits on the membrane, but is free to slide over it, subject to viscous friction. To account for membrane breakup, a more detailed, molecular-level description for lipid bilayers would have been needed (such as in [15, 44]). Generally, the instability of a membrane tube is expected when the hemifission inner lumen radius (ILR) below 2 nm is reached [14]. Fission can be however facilitated by buckling of the tube [45]. It can also be promoted by membrane-bound PH domains whose conformations change within the GTPase cycle [7, 15, 46, 47]. The combination of such effects would probably enable the breakup even at ILRs of 2 − 3 nm.

As has been previously suggested [2, 9], we find that collective motor activity within the dynamin “nanomuscle” resembles that of actomyosin in the muscle (Figure 7). In the muscle, a thick myosin filament slides over a thin filament of actin [30]. Myosin heads bind to actin so that cross-bridges connecting both filaments are formed. As proposed by H. E. Huxley [48], muscle contraction is effectively due to a combination of cyclic conformational changes and a ratchet effect. Depending on its nucleotide state, myosin alternates between open and closed conformations. The transition from the open to the closed states takes place when the bridge exists; it leads to shortening of the bridge and generation of a pulling force. The reverse transition however occurs when the cross-bridge is absent and, therefore, the ratchet is lifted and no force is produced. The muscle is a large assembly where thousands of motors over many filaments operate in parallel.

**Figure 7.**
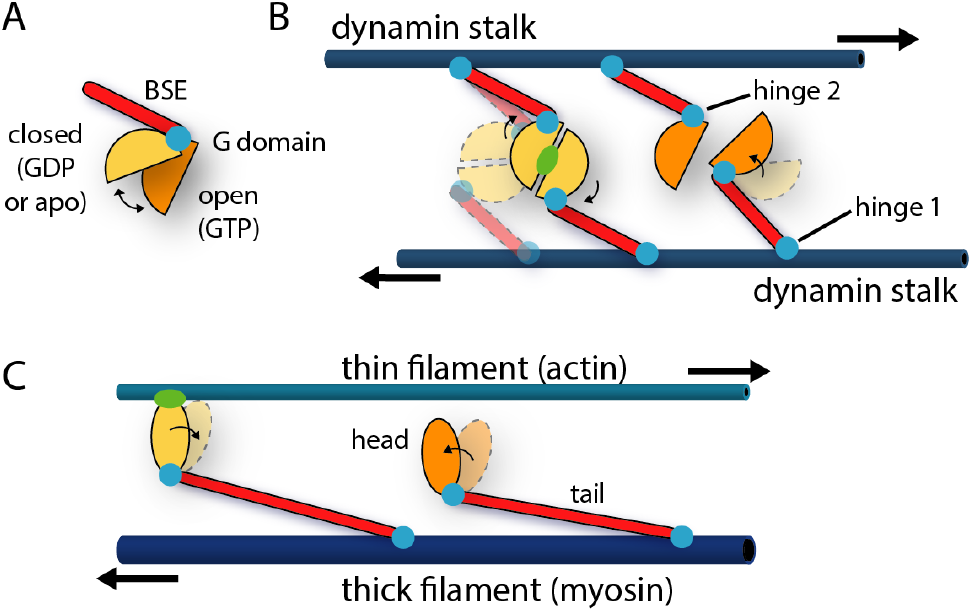
A sliding filaments cross-bridge mechanism for dynamin is suggested by our results. A) smFRET indicates that the motor module exhibits strong nucleotide-dependent conformational shifts in solution. The transitions to closed and open representing, respectively, the power and the recovery strokes in the MM. B,C) Comparison of operation schemes in dynamin and muscle myosin.

For dynamin, in the presence of GTP, cross-bridges between adjacent rungs of a helical stalk filament of dynamin are formed through dimerization of neighboring MMs and preferentially in an open state. A post-hydrolysis shift to the closed state generates a force pulling the filaments with respect to one another, and hence a torque is produced. Binding of GTP and return to the open state can occur only after the disappearance of the cross-bridge. Remarkably, only tens of motor modules are typically involved in this nanomuscle.

In summary, through integrating smFRET measurements, structural modeling, and filament simulations, we have shown that the conformational shifts in the dynamin motor module are strong enough to generate the necessary force to constrict biomembrane tubes when summed over several cooperating cross-bridges. In so doing, we have quantitatively validated the constriction-by-ratchet hypothesis and developed a model that can be generally applied to other membrane-remodeling motors, e.g. those involved in the maintenance and scission of mitochondria.

## 4 Acknowledgements

This work was supported by the European Research Council, ERC-2013-CoG-616024 (O.D), a grant of the German Research Foundation (Collaborative Research Grant 740, From Molecules to Modules) to O.D., by the Japanese Society for Promotion of Science (Grant-in-Aid for Scientific Research (C) 19K03765 to A.M), and through a Humboldt Foundation fellowship to J.N. H.H. thanks the Benoziyo Fund for the Advancement of Science, the Carolito Foundation, the Gurwin Family Fund for Scientific Research, the Leir Charitable Foundation, and the Koshland family. The work was further supported by the Irving and Cherna Moskowitz Center for Nano and Bio-Nano Imaging at the Weizmann Institute of Science. Diffraction data have been collected on BL14.1 at the BESSY II electron storage ring operated by the Helmholtz-Zentrum Berlin. We would particularly like to acknowledge the help and support of Manfred Weiss during the experiment.

## 5 Methods

### 5.1 Cloning, expression, and purification

The pET-M11 vector was used to express a codon-optimized His6-tagged GTPase-BSE (MM) construct (Eurofins, residues 6-322 fused to residues 712-746 with a GSGSGS linker) in BL21 (DE3) Rosetta 2. Cells were grown at 37 °C to an OD600 of 0.4 and the protein was expressed in the presence of 0.15 mM IPTG overnight at 25 °C in TB medium, supplemented with 34 µg/ml of chloramphenicol and 50 µg/ml of kanamycin. Cells were harvested by centrifugation, resuspended in buffer A (20 mM HEPES-NaOH, pH 7.5, 500 mM NaCl, 15 mM imidazole, 5% glycerol) and lysed in a micofluidizer. The cleared lysate after centrifugation was incubated for 30 min with 5 ml Ni-NTA resin at 4 °C on a rocking platform. Afterwards, Ni-NTA beads were washed in batch 3 times with buffer A and transferred into a gravity-flow 20 ml column. Column material was further washed with 10 column volumes (CV) of buffer A supplemented with 2 mM ATP and 5 mM MgCl_2_ and then with 2 CV of buffer A containing 50 mM imidazole. Eventually, the protein was eluted with 5 CV of buffer A containing 250 mM imidazole. The His6 tag was cleaved by incubation with TEV-protease during overnight dialysis against buffer A without imidazole. Subsequently, the protein was re-applied to the Ni-NTA column, collected in the flow-through fraction, and further purified via size-exclusion chromatography on a S200 26/600 Superdex column with running buffer C (20 mM HEPES-NaOH, pH 7.5, 150 mM NaCl). Fractions containing the pure protein were pooled, concentrated to 30-50 mg/ml, aliquoted and shock frozen in liquid nitrogen. The MM wild-type construct as well as various mutant variants were purified following the same protocol. For protein labeling with Alexa fluorophores or for experiments exploring metal-mediated dimerization of the MM constructs, the following mutations were introduced individually or in combination: T165C, R318C, N112C, K142H, N183H. Point mutations in the MM construct were introduced using QuickChange site-directed mutagenesis (Agilent). Cysteine mutations were used for covalent attachment of Alexa Fluor dyes and their locations were chosen to avoid any obvious functional disruption. See Supplementary Information for methods specific to the crystallization and characterization of the Zn^2+^-stabilized dimer.

### 5.2 Protein labeling with fluorescent probes

Thiol-reactive fluorescent dyes (ThermoFisher Scientific) were used for site-specific labeling of cysteines in MM variants. Using AlexaFluor 488 as donor and AlexaFluor 594 as acceptor provides a Föster distance of 5.4 nm [49, 50]. The T165C variant was used to determine the dissociation constants of the artificial dimer and the R318C/N112C double mutant for smFRET measurement. To reduce cysteines, 250 µl of a protein solution at 30 to 40 mg/ml was incubated with 20 mM DTT for 1 h on ice, following DTT removal by gel filtration on a S200 10/300 column equilibrated with buffer F (20 mM HEPES-NaOH, pH 7.5, 5 mM MgCl_2_, 150 mM NaCl, 10 mM KCl). The concentrated proteins were mixed with the corresponding thiol-reactive dye in a 1:2 molar ratio (protein:dye) and incubated for 1 h at room temperature (RT). In case of double labeling by Alexa Fluor 488 and 594, the reaction was started with addition of Alexa Fluor 488 to the protein solution in a 1:0.66 (protein:dye) molar ratio. After 40 min of incubation at RT, Alexa Fluor 594 dye was added in a 1:2 (protein:dye) molar ratio and the reaction was continued for 1 h at RT. The labeling process was terminated by the addition of 10 mM DTT. After 5 min incubation at RT, insoluble material was removed by centrifugation and the sample was applied to a second round of SEC to remove unreacted dye and DTT. Labeling efficacy was calculated according to the manual of Thermo Fisher Scientific; a typical range was 90-100%. All fluorescence-based functional and GTP hydrolysis assays for the monomeric form of the MM constructs were performed in buffer 1 (20 mM HEPES-NaOH, pH 7.5, 150 mM KCl, 4 mM MgCl_2_). Experiments with the artificially dimerized form of the MM-HH variant were performed in buffer 2 (20 mM HEPES-NaOH, pH 7.5, 150 mM KCl, 4 mM MgCl_2_, 160 µM ZnSO_4_). All experiments were carried out at RT if not otherwise stated.

### 5.3 Single-molecule FRET

Single-molecule FRET experiments were performed with a MicroTime 200 confocal system (PicoQuant, Germany) equipped with an Olympus IX73 inverted microscope and two pulsed excitation sources (40 MHz) controlled by a PDL 828-L “Sepia II” (PicoQuant, Germany). Pulsed Interleaved Excitation (PIE) [51] was used to identify molecules with active acceptor and donor dyes. Two light sources, a 485 nm Diode Laser (LDH-D-C-485, PicoQuant) and a white-light laser (Solea, PicoQuant) set to an excitation wavelength of 595 nm were used to excite the donor and the acceptor dyes alternatingly. The laser intensities were adjusted to 100 µW at 485 nm and 30 µW at 595 nm (Pm100D, Thor Labs). The excitation beam was guided through a major dichroic mirror (ZT 470-491/594 rpc, Chroma) to a 60x, 1.2NA water objective (Olympus) that focuses the beam into the sample. The sample was placed in a home made cuvette with a volume of 50 µl (quartz 25 mm diameter round cover slips (Esco Optics), borosilicate glass 6 mm diameter cloning cylinder (Hilgenberg), Norland 61 optical adhesive (Thorlabs)). All measurements were performed at a laser power of 100 µW (485 nm) and 30 µW (585 nm) measured at the back aperture of the objective. Photons emitted from the sample were collected by the same objective and after passing the major dichroic mirror (ZT 470-491/594 rpc, Chroma), the residual excitation light was filtered by a long-pass filter (BLP01-488R, Semrock) and sent through a 100 µm pinhole. The sample fluorescence was detected with two channels. Donor and acceptor fluorescence were separated via a dichroic mirror (T585 LPXR, Chroma) and each color was focused onto a single-photon avalanche diode (SPAD) (Excelitas) with additional bandpass filters: FF03-525/50, (Semrock) for the donor SPAD and FF02-650/100 (Semrock) for the acceptor SPAD. The arrival time of every detected photon was recorded with a HydraHarp 400M time 9 correlated single photon counting module (PicoQuant) and stored with a resolution of 16 ps.

Photons from individual molecules, separated by less than 100 µs were combined into bursts if the total number of photons exceeded 30. Photon counts were corrected for background, acceptor direct excitation, and different detection efficiencies of the individual detectors. A PIE stoichiometry ratio *S <* 0.7 was used to identify molecules with an active acceptor. Identified bursts were corrected for background, differences in quantum yields of donor and acceptor, different collection efficiencies in the detection channels, cross-talk, and direct acceptor excitation. FRET histograms were fitted with a combination of Gaussian functions as previously described.

The labeled MM construct was diluted to a concentration of approximately 50 pM in 20 mM HEPES-NaOH, pH 7.5, 150 mM KCl, 4 mM MgCl_2_ in the absence or presence of increasing amounts of GTP, GTP*γ*S, GMPPCP, GDP. To prevent surface adhesion of the protein and to maximize photon emission, 0.001% Tween 20 (Pierce) was included in the buffer. All measurements were performed at 23 °C.

In the presence of saturating concentrations of GTP and GTP analogs (*>* 1 mM, see Figure S3C), a significant closed population remained, which could not be fully explained by unbound MM (predicted to *<* 2% based on the determined affinities) or GDP contamination. The purity of GTP was 99.4% and of GTP*γ*S was 95%, as determined by HPLC.

### 5.4 Stopped-flow kinetic measurements

All fast kinetic measurements were carried out on Chirascan stopped-flow accessory unit equipped with an independent LED power source (Applied Photophysics). For nucleotide binding studies, fluorescent 2’(3’)-O-(N-methylanthraniloyl) (Mant)-substituted nucleotides were used. The dye fluorescence was excited at 350 nm and the fluorescence change monitored through a Schott 395 nm cut-off filter. In order to measure *k*_on_ values, increasing concentrations of MM construct (2, 3, 4.5, 7, 10.5, 15 and 20 µM) were titrated with 1 µM mant-nucleotide. For each concentration, at least 5 traces originated from the same sample were averaged. A linear regression was performed and the slope provided *k*_on_ with error given by the standard deviation in the slope. For quenching experiments to measure *k*_*rmoff*_, 20 µM protein and 1 µM mant-nucleotide were mixed with 500 µM GTP*γ*S or GDP. Three independent experiments were performed. Rates were determined by single exponential fits and the presented rate represents the average over the three experiments and the error the standard deviation. Error in *K*_d_ is determined by propagating the errors in *k*_on_ and *k*_off_.

In order to detect conformational changes in the MM construct, FRET between Alexa Fluor 488 and 596 fluorescent pairs was recorded. Signals from both dyes were simultaneously detected using two photomultiplier tubes and a Chroma ET525/50m band-pass and Schott 645 nm cut-off filter. G domain/BSE opening was performed by mixing 5 µM of protein and 500 µM of GTP*γ*S. Experiments for measuring G domain/BSE closure were performed as follows: 5 µM protein and 100 µM GTP*γ*S were mixed with 2 mM GDP, where GDP will outcompete GTP*γ*S for binding to the protein. Concentrations of reactants in both syringes of all stopped-flow experiments were two times higher compared to the measurement cell since they were diluted upon mixing in 1:1 ratio. Throughout the manuscript, indicated concentrations are those after mixing.

### 5.5 GTPase assay

GTPase assays were performed in 20 mM HEPES-NaOH, pH 7.5, 150 mM KCl, 4 mM MgCl_2_ containing 5 µM of the MM construct and 1 mM GTP. The reaction carried out at 37 °C. Upon starting the reaction, 2 µl reaction aliquots were withdrawn at different time points and mixed with 2 µl of 1 M HCl to stop the reaction and then diluted with 26 µl of HPLC buffer (100 mM potassium phosphate buffer, pH 6.5, 10 mM tetrabutylammonium bromide, 7.5% acetonitrile). Reaction products were quantified on an HPLC system from Agilent equipped with a Hypersil ODS-2 C18 column (250 × 4 mm). Each time point measurement (5 time points) was done in triplicate with the same sample. Specific hydrolysis rate for each protein concentration was computed by linear regression with error given by the standard deviation of the linear fit. To investigate the effect of Zn^2+^ on artificial dimer activity, 2.5 µM, 10 µM, 40 µM, 80 µM, 160 µM and 400 µM ZnSO_4_ was added to the standard reaction. For testing concentration-dependent GTPase activity, reactions were performed in standard buffer at RT with 5, 15, 30 and 50 µM enzyme. The competition GTPase assay was performed by adding 0.21 or 0.85 mM of GDP to the reaction. See SI Section S1.1.2 for details of the kinetic modeling of the competition assay.

### 5.6 Molecular dynamics (MD) simulations

The conformational dynamics of the MM was studied by means of simplified *structure-based* models (SBMs), a method commonly employed to investigate protein folding. The dual-basin SBM was constructed in such a way that both the known open and closed MM structures were explicit energetic minima of it. The open structure was defined by chain A of the GMPPCP-bound MM (S_O_) [6] and the closed structure by chain A of the GDP-bound MM (S_C_) [22]. The G domains between the two structures are nearly identical. The structural differences mainly reside in the positioning of the first BSE helix relative to the G domain (Figure S5). The weighting *λ* between open and closed in the simulation was determined by matching the simulated open/closed distributions to the smFRET data. See the Supplementary Information for details of the energy potential. Simulations were performed using GROMACS 4.5.3 [52] containing code edits implementing Gaussian contact interactions (available at http://smog-server.org). Single basin simulation topologies were generated using the SMOG2 software [29, 53] with the forcefield “SBM_calpha+gaussian” [54] and combined using an in-house script. The temperature (T=0.92 in reduced units) was chosen such that the average C_*α*_ atom RMSD in the closed MM agreed between a 100 ns explicit solvent simulation using AMBER99SB-ILDN at 310K and the SBM. The temperature was maintained with stochastic dynamics with coupling constant 0.1 and the time step was 0.0005.

To model more closely the distance measured during smFRET, explicit dyes were attached at T165C and R318C. The dye consisted of a linear chain of 6 C_*α*_ beads connected by bonds 3.8 Å in length (like in Figure 2C). The FRET efficiency for each snapshot was calculated based on *E* = (1 + (*d/d*_0_)^6^)^−1^ using the distance *d* between the two terminal dye beads and *d*_0_ = 5.4 nm. To compute the estimated FRET expected in either the open or closed ensemble including thermal fluctuations (Figure 2C) an *λ* of 0.5 (always open) and 1.2 (strongly closed) was used. *λ* of 0.98 and 1.1 matched the smFRET histograms for GTP-bound or GDP-bound, respectively (Figure S5D). Note that the photon counting statistics in the experimental FRET system is sensitive to the timescale of moving between the open and closed states. If the conformational motion time is comparable to the time a single molecule dwells in the laser point (∼1 ms), the FRET distribution will become unimodal. Since separated peaks were observed in the experimental data, the dynamics observed must have been slower than 1 ms. Nonetheless, since averaging always tends to compress the peaks, the simulation data, which was not averaged, was expected to have wider separation between the peaks.

We should note that the experimental smFRET histogram for GTP-bound MM appears to possess an intermediate state (Figure 2A), although a two-state fit with skewed Gaussians is reasonable too. We have not included this intermediate state into the model because there is no crystal structure of it. Although for our purposes here, such an intermediate within GTP-bound dimers would have little effect, only serving to further flatten the already flat *𝓁* distribution.

The flexibility of hinge 1 (Figure S4) was estimated by a C_*α*_ SBM simulation equivalent to that just described except single basin, meaning that any term in the energy function containing the open structure was removed and *λ* = 1. A full monomer (chain A) taken from the apo (closed) crystal structure [18] was used to build the SBM. The PH domain along with stalk domain residues further than two nanometers from hinge 1 were frozen during the simulation, mimicking its filament-bound state. The temperature was the same as above (T=0.92) and the simulation was run for 10^8^ time steps. For analysis of the structural fluctuations in terms of the helical coordinates, the frozen part of the stalk was fit to a stalk domain in the helical reconstruction of Figure 4A. Hinge 1 was identified by PRO322 and hinge 2 was identified by PRO27.

### 5.7 Mean stall forces

The probability distribution for the end-to-end lengths *𝓁* of cross-bridges is determined by the equilibrium distance distribution in pre-hydrolysis GTP-bound dimers, *P* (*𝓁*) = *P*_GTP_(*𝓁*). On the other hand, the contraction force is *F* (*𝓁*) = −*dG*_GDP_(*𝓁*)*/d𝓁* = *k*_*B*_*T* (*d* ln *P*_GDP_(*𝓁*)*/d𝓁*) where *G*_GDP_(*𝓁*) is the Helmholtz free energy of a post-hydrolysis GDP-bound MM dimer. The distance distributions for GTP- and GDP-bound MM dimers are presented in Figure 4C. Thus, the mean force acting along the filament under the stall condition can be determined as

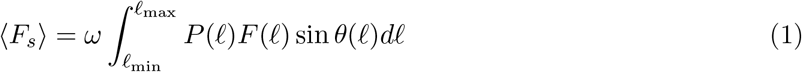

where *θ* is the slant angle and *ω* is the probability (often referred to as the *duty ratio* [30]) to find a MM in the dimerized state. Refer to Figure 4A for the definition of the geometry. The transverse component (*F*_*t*_) is given by the same equation with sin *θ* replaced by − cos *θ*.

The integral in Eq. 1 was computed by using the distribution *P* (*𝓁*) = *A* exp[*G*_GTP_(*𝓁*)*/*k_B_T] and the force *F* (*𝓁*) = −*dG*_GDP_*/d𝓁* with the free energies *G*_GTP_ and *G*_GDP_ obtained in MD simulations, where *A* is a normalization constant for the probability distribution *P* (*𝓁*). In Eq. 1, we have sin *θ* = Δ*s/𝓁* with 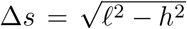. Because of the ratchet property of hinge 1, only configurations with positive displacements Δ*s* should be included when estimating the force. The minimal possible length of a link is *𝓁*_*min*_ = *h*. Functions *P* (*𝓁*) and *F* (*𝓁*) were discretized in 0.5 nm steps and it was assumed that the filament radius is the same in the two rungs.

### 5.8 Elastic treatment of the filament and membrane tube

The code used to implement the constriction model (written in Java) is publicly available at https://bitbucket.org/jk A detailed formulation and analysis of the model for a dynamin filament on a membrane tube, in absence of the motor activity, was given in the previous publication [13]. See the Supplementary Information for a full description. The dynamin filament was represented as an elastic polymer with each bead corresponding to a dynamin dimer. The membrane tube description was based on an axially symmetric continuous Helfrich elastic membrane with stiffness *χ* and under tension *γ*. The membrane stiffness *χ* = 24 k_B_T and the tension *γ* = 0.03 k_B_T/nm^2^ were chosen. At such stiffness, the elastic energy cost of constricting the filament/membrane system is dominated by the membrane energy (Figure S6F). It should be noted that membrane composition and protein insertion can affect the resulting stiffness [55]. As a point of reference a pure POPC membrane has a stiffness around 30 k_B_T. While the true stiffness at the scission neck remains unknown, presumably the known active remodeling of the membrane composition and protein insertions should act to facilitate the scission by shifting the membrane stiffness as low as possible.

### 5.9 Dynamin motors in the constriction model

In the current study, the effects of motor GTPase activity were additionally incorporated into the above-described model. This entailed including explicit cross-bridges representing MM dimers and resolving the kinetics of GTPase cycles. Again, each simulation bead corresponds to a stalk dimer and, thus, has two MMs associated with it. A MM undergoes stochastic transitions between the five ligand states of the GTPase cycle, i.e. Apo and with the nucleotides GDP, GTP, GTP-D or GDP-D (Figure 3). The transitions between three monomeric states are characterized by the nucleotide on/off rates displayed in Figure 1D. The bulk concentrations of GTP and GDP are chosen as 300 µM and 30 µM, respectively (these are typical physiological concentrations). The Apo and GDP states cannot form cross-dimers. Binding of GTP by a MM alters the conformational state from the predominantly closed to the predominantly open at the rate of approximately 300 s^−1^ (Figure 2D,E). Since opening transitions are thus relatively fast, they are not explicitly resolved in the model. Instead, all MMs in the GTP state are assumed to immediately go to the equilibrium open/closed distribution.

Two MM monomers in the GTP state can form the dimeric GTP-D state that represents a weak pre-hydrolysis dimeric state. The GTP state of a monomer transits to a GTP-D state at rate 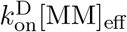, provided that a valid MM partner for dimerization exists. A partner MM is valid if it is also in the GTP state and if the generated cross-dimer would satisfy the conditions *𝓁 < 𝓁*_max_ and Δ*s >* 0, with the latter condition taking into account the ratchet asymmetry of hinge 1. *𝓁*_max_ = 15 nm is the natural extension limit of the dimer as can be seen in the GTP-bound free energy *G*_GDP_(*𝓁*) (Figure 4C). The geometry is explained in Figure S6A. Additionally, due to the excluded volume of the MM dimer, a valid cross-dimer cannot intersect any existing cross-dimers. The broad free energy basin for GTP-bound dimers, found in MD simulations, implies high flexibility (Figure 4C). Therefore, no forces are generated by a GTP-D dimer in the model. The GTP-D dimer dissociates into two GTP MMs at rate 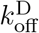.

The transition from a pre-hydrolysis GTP-D dimer to the post-hydrolysis GDP-D dimer takes place at rate *k*_hyd_. Note that hydrolysis is chosen to occur simultaneously in both monomers, i.e. high cooperativity is assumed. Such cooperativity is suggested by our kinetic measurements showing that the hydrolysis is equally fast for the dimers containing either two GTP molecules or one GTP and one GDP (SI Section S1.1.2). The GDP-D dimer generates a force 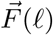 with the magnitude

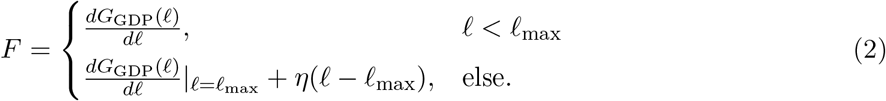

and directed along the vector connecting the two MM attachment points (Figure S6A). The Helmholtz free energy *G*_GDP_(*𝓁*) for the GDP-bound dimer is shown in Figure 4C. As stated above, *𝓁*_max_ = 15 nm. The parameter *η* is set to 3 k_B_T/nm^3^, so that the force steeply increases when *𝓁* exceeds *𝓁*_max_.

The transition from a GTP-D state to the GDP-D state is irreversible due to hydrolysis. The GDP-D state can be left through dissociation into two MM monomers, each in the GDP state, at rate *k*_diss_. When the strain dependence of this rate was taken into account, this was done by choosing

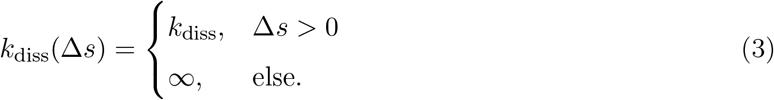

Thus, an oppositely strained dimer dissociates immediately. In the model without strain dependence, *k*_diss_ remains constant regardless of the displacement Δ*s*.

In kinetic simulations involving transitions between the states, the Gillespie algorithm is often used (see, e.g., [56]). In the present study, explicit stochastic simulations of the GTPase cycle were however carried out, because most of the computer time was already needed for integrating the equations of motion for the membrane and the beads. The weighting times of transitions obeyed an exponential distribution with a single rate parameter, i.e. the probability of a transition with rate *k* within time Δ*t* is 1 − exp(−*k*Δ*t*) ≈ *k*Δ*t* for *k*Δ*t* ≪ 1. The transition times (inverse of transition rates) between states were all much longer than the integration time step *δt* of 10 nanoseconds. Therefore, the probability of a transition occurring during a single time step was taken to be *kδt*. The small time step was dictated by the membrane description that had to capture the lipid flows. As a characteristic example, a one-second simulation (i.e., 10^8^ time steps) for a 200-bead filament on a 600 nm long membrane tube required 5 hours on a single core of the Intel Xeon E5-2695 v3 2.30GHz CPU.

The nucleotide association/dissociation rates are given in Figure 1D. Some GTPase-cycle parameters could not be directly measured (Figure 3). Unless otherwise noted, 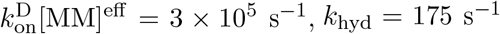, and 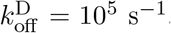. The duty ratio *ω* was varied between 0.37 and 0.94 by changing *k*_diss_ between 100 s^−1^and 3.3 s^−1^. See Section S1.3 for further discussion of the choice of the GTPase cycle kinetic parameters.

## References

[1] Ferguson, S. M & De Camilli, P. (2012) Dynamin, a membrane-remodelling GTPase. Nat. Rev. Mol. Cell Biol. 13, 75–88.

[2] Morlot, S & Roux, A. (2013) Mechanics of dynamin-mediated membrane fission. Annual Review of Biophysics 42, 629–649.

[3] Daumke, O, Roux, A, & Haucke, V. (2014) BAR domain scaffolds in dynamin-mediated membrane fission. Cell 156, 882–892.

[4] Ramachandran, R & Schmid, S. L. (2018) The dynamin superfamily. Current biology: CB 28, R411–R416.

[5] Faelber, K, Gao, S, Held, M, Posor, Y, Haucke, V, Noé, F, & Daumke, O. (2013) Oligomerization of dynamin superfamily proteins in health and disease. Progress in molecular biology and translational science 117, 411–443.

[6] Chappie, J. S, Mears, J. A, Fang, S, Leonard, M, Schmid, S. L, Milligan, R. A, Hinshaw, J. E, & Dyda, F. (2011) A pseudoatomic model of the dynamin polymer identifies a hydrolysis-dependent powerstroke. Cell 147, 209–222.

[7] Kong, L, Sochacki, K. A, Wang, H, Fang, S, Canagarajah, B, Kehr, A. D, Rice, W. J, Strub, M.-P, Taraska, J. W, & Hinshaw, J. E. (2018) Cryo-EM of the dynamin polymer assembled on lipid membrane. Nature pp. 1–17.

[8] Sever, S, Damke, H, & Schmid, S. L. (2000) Garrotes, springs, ratchets, and whips: putting dynamin models to the test. Traffic 1, 385–392.

[9] Antonny, B, Burd, C, De Camilli, P, Chen, E, Daumke, O, Faelber, K, Ford, M, Frolov, V. A, Frost, A, Hinshaw, J. E, Kirchhausen, T, Kozlov, M. M, Lenz, M, Low, H. H, Mcmahon, H, Merrifield, C, Pollard, T. D, Robinson, P. J, Roux, A, & Schmid, S. (2016) Membrane fission by dynamin: what we know and what we need to know. EMBO J. 35, 2270–2284.

[10] Kaksonen M& Roux, A. (2018) Mechanisms of clathrin-mediated endocytosis. Nat. Rev. Mol. Cell Biol. 19, 313–326.

[11] Kozlov, M. M. (2001) Fission of biological membranes: interplay between dynamin and lipids. Traffic 2, 51–65.

[12] McDargh, Z. A, Vázquez-Montejo, P, Guven, J, & Deserno, M. (2016) Constriction by Dynamin: Elasticity versus Adhesion. Biophys. J. 111, 2470–2480.

[13] Noel, J. K, Noé, F, Daumke, O, & Mikhailov, A. S. (2019) Polymer-like Model to Study the Dynamics of Dynamin Filaments on Deformable Membrane Tubes. Biophys. J. 117, 1870–1891.

[14] Kozlovsky, Y & Kozlov, M. M. (2003) Membrane fission: model for intermediate structures. Biophys. J. 85, 85–96.

[15] Pannuzzo, M, McDargh, Z. A, & Deserno, M. (2018) The role of scaffold reshaping and disassembly in dynamin driven membrane fission. eLife 7, 2270.

[16] Lenz, M, Prost, J, & Joanny, J.-F. (2008) Mechanochemical action of the dynamin protein. Phys. Rev. E 78, 011911.

[17] Morlot, S, Lenz, M, Prost, J, Joanny, J.-F, & Roux, A. (2010) Deformation of dynamin helices damped by membrane friction. Biophys. J. 99, 3580–3588.

[18] Faelber, K, Posor, Y, Gao, S, Held, M, Roske, Y, Schulze, D, Haucke, V, Noé, F, & Daumke, O. (2011) Crystal structure of nucleotide-free dynamin. Nature 477, 556–560.

[19] Ford, M. G. J, Jenni, S, & Nunnari, J. (2011) The crystal structure of dynamin. Nature 477, 561–566.

[20] Chappie, J. S, Acharya, S, Liu, Y.-W, Leonard, M, Pucadyil, T. J, & Schmid, S. L. (2009) An intramolecular signaling element that modulates dynamin function in vitro and in vivo. Mol. Biol. Cell 20, 3561–3571.

[21] Chappie, J. S, Acharya, S, Leonard, M, Schmid, S. L, & Dyda, F. (2010) G domain dimerization controls dynamin’s assembly-stimulated GTPase activity. Nature 465, 435–440.

[22] Anand, R, Eschenburg, S, & Reubold, T. F. (2016) Crystal structure of the GTPase domain and the bundle signalling element of dynamin in the GDP state. Biochemical and Biophysical Research Communications 469, 76–80.

[23] Rennie, M. L, McKelvie, S. A, Bulloch, E. M. M, & Kingston, R. L. (2014) Transient dimerization of human MxA promotes GTP hydrolysis, resulting in a mechanical power stroke. Structure (London, England: 1993) 22, 1433–1445.

[24] Chen, Y, Zhang, L, Graf, L, Yu, B, Liu, Y, Kochs, G, Zhao, Y, & Gao, S. (2017) Conformational dynamics of dynamin-like MxA revealed by single-molecule FRET. Nat. Commun. 8, 15744.

[25] Traut, T. W. (1994) Physiological concentrations of purines and pyrimidines. Molecular and cellular biochemistry 140, 1–22.

[26] Binns, D. D, Barylko, B, Grichine, N, Atkinson, M. A, Helms, M. K, Jameson, D. M, Eccleston, J. F, & Albanesi, J. P. (1999) Correlation between self-association modes and GTPase activation of dynamin. Journal of protein chemistry 18, 277–290.

[27] Binns, D. D, Helms, M. K, Barylko, B, Davis, C. T, Jameson, D. M, Albanesi, J. P, & Eccleston, J. F. (2000) The mechanism of GTP hydrolysis by dynamin II: a transient kinetic study. Biochemistry 39, 7188–7196.

[28] Chen, Y.-J, Zhang, P, Egelman, E. H, & Hinshaw, J. E. (2004) The stalk region of dynamin drives the constriction of dynamin tubes. Nat. Struct. Mol. Biol. 11, 574–575.

[29] Noel, J. K, Levi, M, Raghunathan, M, Lammert, H, Hayes, R. L, Onuchic, J. N, & Whitford, P. C. (2016) SMOG 2: A Versatile Software Package for Generating Structure-Based Models. PLOS Comput. Biol. 12, e1004794.

[30] Sweeney H. L & Houdusse, A. (2010) Structural and functional insights into the Myosin motor mechanism. Annual Review of Biophysics 39, 539–557.

[31] Cocucci, E, Gaudin, R, & Kirchhausen, T. (2014) Dynamin recruitment and membrane scission at the neck of a clathrin-coated pit. Mol. Biol. Cell 25, 3595–3609.

[32] Veigel, C, Schmitz, S, Wang, F, & Sellers, J. R. (2005) Load-dependent kinetics of myosin-V can explain its high processivity. Nat. Cell Biol. 7, 861–869.

[33] Oguchi, Y, Mikhailenko, S. V, Ohki, T, Olivares, A. O, De La Cruz, E. M, & Ishiwata, S. (2008) Load-dependent ADP binding to myosins V and VI: implications for subunit coordination and function. Proc. Nat. Acad. Sci. USA 105, 7714–7719.

[34] Iwaki, M, Iwane, A. H, Shimokawa, T, Cooke, R, & Yanagida, T. (2009) Brownian search-and-catch mechanism for myosin-VI steps. Nat. Chem. Biol. 5, 403–405.

[35] Kaya, M, Tani, Y, Washio, T, Hisada, T, & Higuchi, H. (2017) Coordinated force generation of skeletal myosins in myofilaments through motor coupling. Nat. Commun. 8, 16036–16013.

[36] Watanabe, S & Boucrot, E. (2017) Fast and ultrafast endocytosis. Current opinion in cell biology 47, 64–71.

[37] Morlot, S, Galli, V, Klein, M, Chiaruttini, N, Manzi, J, Humbert, F, Dinis, L, Lenz, M, Cappello, G, & Roux, A. (2012) Membrane shape at the edge of the dynamin helix sets location and duration of the fission reaction. Cell 151, 619–629.

[38] Takeda, T, Kozai, T, Yang, H, Ishikuro, D, Seyama, K, Kumagai, Y, Abe, T, Yamada, H, Uchihashi, T, Ando, T, & Takei, K. (2018) Dynamic clustering of dynamin-amphiphysin helices regulates membrane constriction and fission coupled with GTP hydrolysis. eLife 7, 393.

[39] Roux, A, Uyhazi, K, Frost, A, & De Camilli, P. (2006) GTP-dependent twisting of dynamin implicates constriction and tension in membrane fission. Nature 441, 528–531.

[40] Obar, R. A, Collins, C. A, Hammarback, J. A, Shpetner, H. S, & Vallee, R. B. (1990) Molecular cloning of the microtubule-associated mechanochemical enzyme dynamin reveals homology with a new family of GTP-binding proteins. Nature 347, 256–261.

[41] Damke, H, Baba, T, Warnock, D. E, & Schmid, S. L. (1994) Induction of mutant dynamin specifically blocks endocytic coated vesicle formation. Journal of Cell Biology 127, 915–934.

[42] Gao, S, von der Malsburg, A, Dick, A, Faelber, K, Schröder, G. F, Haller, O, Kochs, G, & Daumke, O. (2011) Structure of myxovirus resistance protein a reveals intra-and intermolecular domain interactions required for the antiviral function. Immunity 35, 514–525.

[43] Kadosh, A, Colom, A, Yellin, B, Roux, A, & Shemesh, T. (2019) The tilted helix model of dynamin oligomers. Proc. Nat. Acad. Sci. USA 116, 12845–12850.

[44] Huang, M.-J, Kapral, R, Mikhailov, A. S, & Chen, H.-Y. (2012) Coarse-grain model for lipid bilayer self-assembly and dynamics: multiparticle collision description of the solvent. J. Chem. Phys. 137, 055101.

[45] Vasan, R, Rudraraju, S, Akamatsu, M, Garikipati, K, & Rangamani, P. (2019) A mechanical model reveals that non-axisymmetric buckling lowers the energy barrier associated with membrane neck constriction. Soft Matter.

[46] Shnyrova, A. V, Bashkirov, P. V, Akimov, S. A, Pucadyil, T. J, Zimmerberg, J, Schmid, S. L, & Frolov, V. A. (2013) Geometric catalysis of membrane fission driven by flexible dynamin rings. Science 339, 1433–1436.

[47] Mattila, J.-P, Shnyrova, A. V, Sundborger, A. C, Hortelano, E. R, Fuhrmans, M, Neumann, S, Müller, M, Hinshaw, J. E, Schmid, S. L, & Frolov, V. A. (2015) A hemi-fission intermediate links two mechanistically distinct stages of membrane fission. Nature 524, 109–113.

[48] Huxley, H. E. (1969) The mechanism of muscular contraction. Science 164, 1356–1365.

[49] Schuler, B, Lipman, E. A, & Eaton, W. A. (2002) Probing the free-energy surface for protein folding with single-molecule fluorescence spectroscopy. Nature 419, 743–747.

[50] Schuler, B & Hofmann, H. (2013) Single-molecule spectroscopy of protein folding dynamics– expanding scope and timescales. Curr. Opin. Struct. Biol. 23, 36–47.

[51] Müller, B. K, Zaychikov, E, Bräuchle, C, & Lamb, D. C. (2005) Pulsed interleaved excitation. Biophys. J. 89, 3508–3522.

[52] Pronk, S, Páll, S, Schulz, R, Larsson, P, Bjelkmar, P, Apostolov, R, Shirts, M. R, Smith, J. C, Kasson, P. M, van der Spoel, D, Hess, B, & Lindahl, E. (2013) GROMACS 4.5: a high-throughput and highly parallel open source molecular simulation toolkit. Bioinformatics 29, 845–854.

[53] Noel, J. K, Whitford, P. C, & Onuchic, J. N. (2012) The Shadow Map: A General Contact Definition for Capturing the Dynamics of Biomolecular Folding and Function. J. Phys. Chem. B 116, 8692–8702.

[54] Lammert, H, Schug, A, & Onuchic, J. N. (2009) Robustness and generalization of structure-based models for protein folding and function. Proteins: Struct., Funct., Bioinf. 77, 881–891.

[55] Dimova, R. (2014) Recent developments in the field of bending rigidity measurements on membranes. acis 208, 225–234.

[56] Kubo, S, Niina, T, & Takada, S. (2020) Molecular dynamics simulation of proton-transfer coupled rotations in ATP synthase FO motor. Scientific reports 10, 8225–8216.

