## Supplementary Information for "Quantification and demonstration of the constriction-by-rachet mechanism in the dynamin molecular motor"

### Supplementary Information for “Title XXX”

### S1 Supplementary Text and Figures

#### S1.1 Dynamin G domain dimerization and GTPase kinetics

##### S1.1.1 Linear dependence of the specific hydrolysis rate

The observed assembly-based activation of GTPase activity in dynamin is thought to be driven (i) by the filament structure aligning the G domains for dimerization, and (ii) by the increase of the GTPase rate for the dimers formed via their G interface [1]. If the dimers enhance the GTPase rate, then the GTPase activity during the *in vitro* study of our MM construct should strongly depend on the concentration of MM dimers. At saturating GTP concentration, the GDP production rate $k_{\mathrm{hyd}}^{\mathrm{total}}$ is given by

$$k_{\mathrm{hyd}}^{\mathrm{total}} = 2k_{\mathrm{hyd}}[\mathrm{TMMTMM}] + k_{\mathrm{hyd}}^{\mathrm{basal}}[\mathrm{TMM}] \tag{S1}$$

where $k_{\mathrm{hyd}}$ is the hydrolysis rate of a GTP-bound dimer, [TMMTMM] and [TMM] are the concentrations of GTP-bound (T) MM dimers and monomers, and the factor 2 stems from the fact that the dimer binds two GTP molecules. The kinetics of MM dimerization, again under saturating GTP concentrations, is given by

$$\frac{d[\mathrm{TMMTMM}]}{dt} = k_{\mathrm{on}}^{\mathrm{D}}[\mathrm{TMM}][\mathrm{TMM}] - k_{\mathrm{off}}^{\mathrm{D}}[\mathrm{TMMTMM}] - k_{\mathrm{hyd}}[\mathrm{TMMTMM}]. \tag{S2}$$

At the steady state with ($d[\mathrm{TMMTMM}]/dt{=}0$) and for a small fraction of dimers ($[\mathrm{TMMTMM}] \ll [\mathrm{TMM}]$), Eqs. S1 and S2 yield a specific hydrolysis rate $k_{\mathrm{hyd}}^{\mathrm{spec}}$, i.e. the observed rate of hydrolysis per GTPase molecule, given by

$$k_{\mathrm{hyd}}^{\mathrm{spec}}([M]) = k_{\mathrm{hyd}}^{\mathrm{total}}/[\mathrm{TMM}] \approx \alpha[\mathrm{TMM}] + k_{\mathrm{hyd}}^{\mathrm{basal}} \tag{S3}$$

that is linear with respect to the MM concentration, with an intercept given by the basal rate and the slope $\alpha$ given by $\alpha = 2k_{\mathrm{hyd}}k_{\mathrm{on}}^{\mathrm{D}}/(k_{\mathrm{hyd}} + k_{\mathrm{off}}^{\mathrm{D}})$. With the simpler MM construct, the linear result shown in Figure 1F confirms that the GTPase activity is activated by dimerization and the slope gives $\alpha = 3.5\ \mathrm{s}^{-1}\mathrm{mM}^{-1}$. Note that Eq. S3 breaks down when $[\mathrm{TMM}] \sim K_{\mathrm{d}}^{\mathrm{dimer}}$, implying that $K_{\mathrm{d}}^{\mathrm{dimer}} \gg 50\ \mu\mathrm{M}$ (Figure 1F).

Below, we discuss two situations leading to more complicated GTPase dynamics than Eq. S3. In the first of them, the assumption of the saturating GTP concentration is dropped in order to analyze competition between GTP and GDP. In the second one, the assumption that the population of MM dimers is much smaller than that of MM monomers is dropped to consider GTPase dynamics in an assembled filament.

##### S1.1.2 GTPase competition assay

Running the GTPase reaction until all GTP is consumed provides a richer dynamics to extract kinetic parameters since GDP can compete for the active site. Thus, the GTPase rate will depend on the relative affinity of GTP versus GDP for the nucleotide-binding pocket, and parameter fitting of a theoretical reaction scheme to the experimental kinetic data provides an estimate of the GTP binding rate $k_{\text{on}}^{\text{GTP}}$ (useful since mant-GTP binding showed complicated kinetics).

The competition assay involved mixing MM, GTP, and GDP and running the reaction beyond the linear regime. The concentration of GDP was monitored as a function of time. The kinetic model is given by:

$$\begin{aligned}
\frac{d[\text{MM}]}{dt} &= -k_{\text{on}}^{\text{GTP}}[\text{T}][\text{MM}] + k_{\text{off}}^{\text{GTP}}[\text{TMM}] - k_{\text{on}}^{\text{GDP}}[\text{D}][\text{MM}] + k_{\text{off}}^{\text{GDP}}[\text{DMM}] \\
\frac{d[\text{TMM}]}{dt} &= k_{\text{on}}^{\text{GTP}}[\text{T}][\text{MM}] - k_{\text{off}}^{\text{GTP}}[\text{TMM}] \\
\frac{d[\text{DMM}]}{dt} &= k_{\text{on}}^{\text{GDP}}[\text{D}][\text{MM}] - k_{\text{off}}^{\text{GDP}}[\text{DMM}] \quad \leftarrow \text{Competitive inhibition} \\
\frac{d[\text{T}]}{dt} &= -k_{\text{on}}^{\text{GTP}}[\text{T}][\text{MM}] + k_{\text{off}}^{\text{GTP}}[\text{TMM}] - \alpha[\text{TMM}][\text{TMM}] - \alpha^*[\text{TMM}][\text{DMM}] - k_{\text{hyd}}^{\text{basal}}[\text{TMM}] \\
\frac{d[\text{D}]}{dt} &= -k_{\text{on}}^{\text{GDP}}[\text{D}][\text{MM}] + k_{\text{off}}^{\text{GDP}}[\text{DMM}] + \alpha[\text{TMM}][\text{TMM}] + \alpha^*[\text{TMM}][\text{DMM}] + k_{\text{hyd}}^{\text{basal}}[\text{TMM}] \qquad \text{(S4)}
\end{aligned}$$

Here, [MM] is the concentration of motor domains, [TMM] and [DMM] are those of the motor domains bound to GTP or GDP, [T] and [D] are the concentrations of GTP and GDP. Because $K_{\text{d}}^{\text{dimer}} \gg 50$ μMand the employed protein concentrations are less than 50 μM, the population fraction of the dimers ([TMMTMM]) is negligibly small and, thus, does not need to be explicitly tracked. $k_{\text{hyd}}^{\text{basal}}$ is the monomer hydrolysis rate. If only $k_{\text{hyd}}^{\text{basal}}$ were present (i.e. $\alpha = \alpha^* = 0$), this would have been the well-known reaction scheme for a catalytic enzyme with a competitive inhibitor.

The kinetic model was fit to the data shown in Figures S1C-D, except for the run with 50 μM protein because its hydrolysis rate was too high to be accurately measured. The following parameters from the stopped-flow experiments were used: $k_{\text{off}}^{\text{GTP}} = 108$ s$^{-1}$, $k_{\text{on}}^{\text{GDP}} = 7.2$ s$^{-1}$μM$^{-1}$, and $k_{\text{off}}^{\text{GDP}} = 120$ s$^{-1}$. Note that these values were determined with mant-labeled nucleotides. Additionally, from the GTPase data in the linear regime, we know $\alpha = 3.5$ s$^{-1}$μM $^{-1}$ (Figure 1F). The result of fitting the GTP binding rate, while holding $\alpha = 3.5$ s$^{-1}$μM $^{-1}$ and $\alpha^* = 0$, is shown in Figure S1E. Although the fit is roughly satisfactory, the model GDP production clearly lags behind the measured GDP production for the higher GDP concentrations. Allowing both $k_{\text{on}}^{\text{GTP}}$ and $\alpha$ to be optimized yields $\alpha \neq 3.5$ s$^{-1}$μM $^{-1}$. In order to obtain a quantitative fit, an additional kinetic process needed to be introduced, i.e. the hydrolysis in the dimers formed between a GTP-bound monomer and a GDP-bound monomer, so that $\alpha^* > 0$ (Figure S1F). Interestingly, when $k_{\text{on}}^{\text{GTP}}$, $\alpha$ and $\alpha^*$ are all taken as free parameters, $\alpha = 3.5$ s$^{-1}$μM $^{-1}$ is recovered. This increases the confidence in the model since $\alpha = 3.5$ s$^{-1}$μM $^{-1}$ was clearly obtained in the linear regime.

The fact that $\alpha \approx 2\alpha^*$ (Table S1) provides support for the assumption, made in the constriction model, that hydrolysis occurs simultaneously in both G domains within an MM dimer. Although it may occur not exactly simultaneously, this result suggests that the hydrolysis in one domain does not cripple the hydrolysis in its partner. Therefore, if the hydrolysis tends to stabilize the dimer, the other monomer would rapidly hydrolyze too.

The fitting was performed by using the `minpack.lm` function `nls.lm` in the R language. This function minimizes, by employing the Levenberg-Marquart algorithm, the root-mean-squared differences between the fit and the experimental data summed over all experimental points. Since this method can be sensitive to initial conditions, a range of initial parameter values was taken. Only the results with the lowest mean-squared error are shown in Table S2. The uncertainties in the four-parameter fit are estimated by randomly adding [-0.007,0.007] to each experimental point and rerunning the non-linear parameter optimization using the fitted values as the initial conditions each time (0.007 was the rms error between the fit and the experimental points). The standard deviation of each parameter over 100 trials is presented as its uncertainty in Table S2.

##### S1.1.3 Specific hydrolysis rate within the filament (and in constriction simulations)

The GTPase activity within the filament is more complicated than described by Eq. S3 because the assumption that the concentration of GTP-bound MM monomers ([TMM]) is much larger than that of the GTP-bound MM dimers ([TMMTMM]) might break down within it. The dimerization can be strongly driven by a large effective concentration of GTP-bound MMs within the filament structure, which we denote as $[\text{MM}]^{\text{eff}}$. The lower limit for $[\text{MM}]^{\text{eff}}$ is obtained by considering the geometry of the filament and noting that the molarity of one MM in

| # free param | $k_{\text{on}}^{\text{GTP}}$ s$^{-1}$µM $^{-1}$ | $\alpha$ s$^{-1}$µM $^{-1}$ | $\alpha^*$ s$^{-1}$µM $^{-1}$ | $k_{\text{hyd}}^{\text{basal}}$ s$^{-1}$ | ms-error |
|---|---|---|---|---|---|
| 1 | 12 | **3.5** | **0** | **0** | 0.014 |
| 2 | 16 | 2.8 | **0** | **0** | 0.013 |
| 2 | 6.9 | **3.5** | 1.3 | **0** | 0.006 |
| 3 | 6.8 | 3.5 | 1.3 | **0** | 0.007 |
| 4 | 6.6 ±0.8 | 3.5 ± 0.3 | 1.3 ± 0.2 | 0.01 ± 0.007 | 0.007 |

Table S1: Parameters extracted with a non-linear least-squares fit between the kinetic model predictions and the experimental data at every time point (41 in total). Final mean-squared (ms) errors are shown. Estimates of the uncertainty of the fitted parameters are only calculated for the four-parameter fit. Bolded entries were frozen at the indicated values during fitting.

a box of volume $5.6 \times 10 \times 10$ nm$^3$ is 3 mM. Since the MMs are attached by BSE to the filament rather than freely diffusing within this box, and, additionally, they tend to point towards their neighbors, $[\text{MM}]^{\text{eff}}$ must be larger than 3 mM. At such high concentrations, it may well be that the equilibrium is shifted to the dimers.

In the case where [TMM] $\sim$ [TMMTMM], the steady-state approximation can still be used for $k_{\text{hyd}}^{\text{spec}}$. Given an effective local concentration $[\text{MM}]^{\text{eff}}$ of the monomers, the MM population partitions into monomers and dimers, i.e. $[\text{MM}]^{\text{eff}} = [\text{TMM}]+2[\text{TMMTMM}]$. Note that [TMM] is now given by the steady-state approximation of Eq. S3 without assuming [TMM] $\gg$ [TMMTMM]. It yields

$$[\text{TMM}] = \frac{-(k_{\text{off}}^{\text{D}} + k_{\text{hyd}})/2 + \sqrt{(k_{\text{off}}^{\text{D}} + k_{\text{hyd}})^2/4 + 2k_{\text{on}}^{\text{D}}(k_{\text{off}}^{\text{D}} + k_{\text{hyd}})[\text{MM}]^{\text{eff}}}}{2k_{\text{on}}^{\text{D}}}$$
$$= \frac{k_{\text{off}}^{\text{D}} + k_{\text{hyd}}}{4k_{\text{on}}^{\text{D}}}\left(\sqrt{1 + \frac{8k_{\text{on}}^{\text{D}}[\text{MM}]^{\text{eff}}}{k_{\text{off}}^{\text{D}} + k_{\text{hyd}}}} - 1\right) \tag{S5}$$

This steady-state value for [MM] can be further used to find

$$k_{\text{hyd}}^{\text{spec}} = \frac{2k_{\text{hyd}}[\text{TMMTMM}]}{[\text{MM}]^{\text{eff}}} = k_{\text{hyd}}\left(\frac{[\text{MM}]^{\text{eff}} - [\text{TMM}]}{[\text{MM}]^{\text{eff}}}\right) \tag{S6}$$

When employed in our polymer model, the rate constant $k_{\text{hyd}}^{\text{spec}}$ is defined as above in terms of the three dimer rates $k_{\text{hyd}}$, $k_{\text{on}}^{\text{D}}$, and $k_{\text{off}}^{\text{D}}$, and the effective concentration $[\text{MM}]^{\text{eff}}$. The dependence given by Eq. S6 can be compared to the relationship $k_{\text{hyd}}^{\text{spec}} = \alpha[\text{TMM}]$ (Eq. S3) in the GTPase experiments at low protein concentrations (Figure 1E), where

$$\alpha = \frac{2k_{\text{hyd}}k_{\text{on}}^{\text{D}}}{k_{\text{hyd}} + k_{\text{off}}^{\text{D}}} = 3.5\ \text{s}^{-1}\text{mM}^{-1}. \tag{S7}$$

While $k_{\text{hyd}}^{\text{spec}}$ sets the maximum possible (average) hydrolysis rate, note that the observed (average) hydrolysis rate in the simulations will always be lower, since partner MMs are not always available for dimerization.

#### S1.2 Design and analysis of the MM dimer simulations

As stated in the main text, the smFRET experiments were limited to MM monomers, and thus, the distance distributions within the MM dimer were instead studied using molecular simulations. The dimer was created by connecting two MM monomers using the interactions within the G interface (Figure 4B). Individually, the simulated MM monomers were tuned to reproduce the experimental smFRET distribution (i.e. ratio between open and closed states). In this way, reaction coordinates within the dimer (namely the end-to-end distance $\ell$) could be studied within an ensemble that is consistent with the experimental data on the monomer. This design, which anchored the simulations to the experimental data, calibrated the units of the simulated free energies.

The fact that the energetics were constrained by experimental data allowed us to employ a simplified protein model called a *structure-based* model (SBM). SBMs are commonly used in the study of protein folding and dynamics [2, 3]. They are known as structure-based because the global minimum of the energy function is assigned based on a crystallographic structure. In them, electrostatics and solvation effects are implicitly accounted for, since stabilizing interactions in a model describe effective residue interactions after averaging over numerous interactions that stabilize a particular conformational crystal state. Here, SBMs are particularly well-suited because the open and closed MM configurations were both previously crystallized (Figure 2B). As a self-consistency check, (single-basin) structure-based MD simulations of the monomeric MM based on either the GMPPCP crystal structure (open) or on the GDP crystal structure (closed) yielded FRET levels for the open and closed states that were similar to those observed in smFRET (Figure 2C).

The essential feature of the MM monomer is that it can transition between open and closed states. The simplified energetics of SBMs allowed a straightforward matching to the experimentally-measured distributions. A dual-basin SBM was constructed in such a way that both the open and closed MM structures were explicit energetic minima of it (see Methods). The specific construction of the dual-basin SBM is unlikely to influence the presented predictions because the experimental FRET information was used to calibrate the energetics. The principal results were (i) the length dependence of the force for GTP-bound and GDP-bound MM dimers and (ii) validation of open and closed crystal structures for describing smFRET data. They refer to geometrical features not sensitive to potential's details.

Using the dual-basin SBM, the free energy profile with respect to the FRET coordinate of the simulated GTP-bound MM monomer had two minima, corresponding to the open and closed states, whereas the GDP-bound MM monomer showed a single minima at the closed state (Figure S5C). Connecting two monomers via the G interface to create the MM dimer did not introduce correlations between the monomers as can be seen in the two-dimensional free energy profiles (Figure S5F). That is, each monomer independently samples open and closed states.

Matching to the experimental distributions amounted to varying a single parameter $\lambda$ that controlled the relative weighting of the open to closed states. Using the definition that the open states have FRET <0.5 and the closed states have FRET >0.5, $\lambda$ was chosen in such a way that total population ratios of the open and closed states were the same in MD simulations and smFRET (Figure S5D). Such rather coarse matching was used because the detailed FRET and simulation histograms are not expected to completely agree, because of statistical averaging at the experimental level and due to the simplifications made in MD simulations (Figure S5D).

The MM dimer simulations allowed us to determine free energies $G_{\mathrm{GTP}}$ and $G_{\mathrm{GDP}}$ in different nucleotide states as functions of the dimer length $\ell$ (Figure S5E). Remarkably, we find that, in the GTP-bound state, the free energy has a single, broad and shallow, minimum resulting in a wide and flat equilibrium distribution for $\ell$. On the other hand, two-variable distributions in the presence of GTP (Figure S5F) showed that the MM dimer had open-open, open-closed, and closed-closed conformations that corresponded to various combinations of the open or closed conformations of the two monomers that make it up. Therefore, the appearance of a single broad energy minimum in $G_{\mathrm{GTP}}(\ell)$ is a result of projecting a two-dimensional energy landscape onto a single coordinate. Such simplification of energy landscapes when projected on a single distance-based reaction coordinate is known, e.g., to arise in single-molecule pulling experiments [4]. For the GDP-bound case, the determined dependence of free energy $G_{\mathrm{GDP}}$ on variable $\ell$ mirrors the monomer distributions which are strongly biased towards the closed state.

#### S1.3 Parameters used in membrane constriction simulations

The most important parameter is the duty ratio $\omega$ that specifies during what fraction of the MM GTPase cycle a force is produced, i.e. the MM is found in the strongly-dimerized cross-bridge state. In terms of the GTPase cycle kinetics (Figure 2D), the duty ratio can be expressed as

$$\omega = (k_{\mathrm{diss}} T)^{-1} \tag{S8}$$

where $T = 1/k_{\mathrm{hyd}}^{\mathrm{spec}} + 1/k_{\mathrm{diss}} + 1/k_{\mathrm{N}}$ is the average cycle time, $k_{\mathrm{N}}$ is the nucleotide exchange rate (GDP to GTP) and $k_{\mathrm{diss}}$ is the dimer dissociation rate. We know $k_{\mathrm{N}} \approx 120\ \mathrm{s}^{-1}$, the GDP dissociation rate. Since the duty ratio however also depends on experimentally inaccessible rates, i.e. on the hydrolysis rate $k_{\mathrm{hyd}}^{\mathrm{spec}}$ within the constrained environment of the filament and on the dissociation rate $k_{\mathrm{diss}}$, simulations over a series of dissociation rates $k_{\mathrm{diss}}$ from 3.3 $\mathrm{s}^{-1}$ to 100 $\mathrm{s}^{-1}$ and hydrolysis rates $k_{\mathrm{hyd}}^{\mathrm{spec}}$ from 10 $\mathrm{s}^{-1}$ to 120 $\mathrm{s}^{-1}$, yielding duty ratios $\omega$ varying between 0 and 1, were performed. It was found that the steady-state constriction radius is almost entirely dependent on the duty ratio $\omega$, with only weak dependence observed under the variation of the individual rates (i.e. $k_{\mathrm{diss}}$, $k_{\mathrm{hyd}}$, $k_{\mathrm{on}}^{\mathrm{D}}$, $k_{\mathrm{off}}^{\mathrm{D}}$) at constant $\omega$.

Due to the low affinity of MM dimers, rate constants $k_{\mathrm{hyd}}$, $k_{\mathrm{on}}^{\mathrm{D}}$, and $k_{\mathrm{off}}^{\mathrm{D}}$ could not be determined directly from the experiments. Additionally, the effective concentration $[\mathrm{MM}]^{\mathrm{eff}}$ of MMs in the filament is unknown. Because a combination of these four parameters yields $\alpha$, one of them can be determined, but three other parameters still remain arbitrary (see Section S1.1.3 for discussion). In the reported simulations, we have chosen a regime where the rate $k_{\mathrm{hyd}}^{\mathrm{spec}}$ was not limiting, it was equal to $k_{\mathrm{hyd}}^{\mathrm{spec}} = 120\ \mathrm{s}^{-1}$, the same as that of GDP dissociation. Specifically, the parameters were $[\mathrm{MM}]^{\mathrm{eff}} = 300$ mM, $k_{\mathrm{on}}^{\mathrm{D}} = 1\ \mathrm{s}^{-1}\mu\mathrm{M}^{-1}$ (a typical value for protein association rates [5]), $k_{\mathrm{off}}^{\mathrm{D}} = 10^5\ \mathrm{s}^{-1}$ (or equivalently $K_{\mathrm{d}}^{\mathrm{dimer}} = 100$ mM). This sets $k_{\mathrm{hyd}} = 175\ \mathrm{s}^{-1}$ based on $\alpha$. For the case of the strain-dependent dissociation rate $k_{\mathrm{diss}}$, the constriction speed becomes limited by $k_{\mathrm{hyd}}^{\mathrm{spec}}$. In absence of the strain dependence, the constriction speed is instead mostly controlled by $k_{\mathrm{diss}}$ and only weakly dependent on $k_{\mathrm{hyd}}^{\mathrm{spec}}$ (since large $\omega$ requires $k_{\mathrm{diss}} \ll k_{\mathrm{hyd}}^{\mathrm{spec}}$). For the same reason, at large duty ratios $\omega$, the steady-state ILR is only weakly dependent on $k_{\mathrm{hyd}}^{\mathrm{spec}}$ (Eq. S8). Note that GTPase-activation experiments imply a lower bound of $4-6\ \mathrm{s}^{-1}$ for the overall turnover rate [6].

The results of simulations where $k_{\mathrm{hyd}}^{\mathrm{spec}}$ larger than $k_{\mathrm{diss}}$ are not displayed in the figures. This regime necessarily has low duty ratios (Eq. S8), and thus, tight constriction could not be observed.

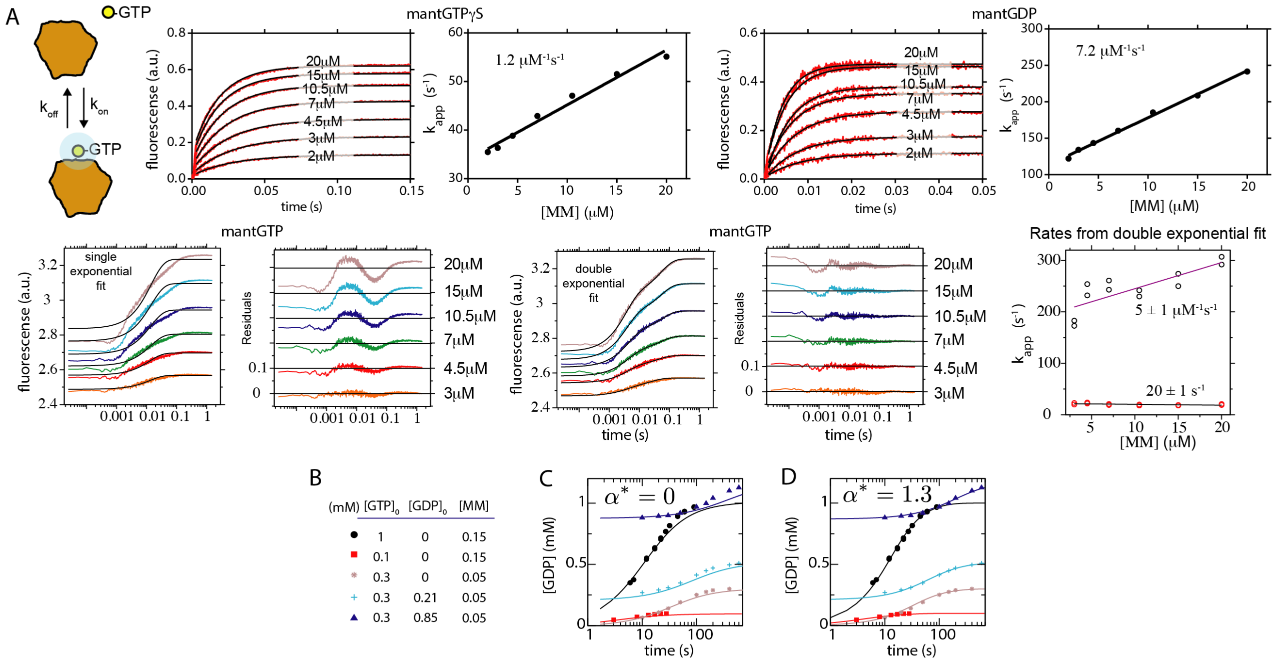

Figure S1: Nucleotide and GTPase kinetics. A) Nucleotide binding to the MM construct measured with mant-labelled nucleotides and stopped flow. The fluorescence of the mant dye, which is sensitive to a change of environment from solvent to protein, was monitored by stopped-flow. 1 μM of mant-labeled nucleotide was mixed with protein concentrations between 2 μM and 20 μM and $k_{\text{app}}$ was determined by an exponential fit. A linear fit of $k_{\text{app}}$ versus protein concentration then provided $k_{\text{app}}$ (Top) Single exponential fits to the fluorescence change for mant-GTP$\gamma$S and mant-GDP. (Bottom) Single and double exponential fits to the fluorescence change for mant-GTP. Double exponential fits more closely to the data. One rate (5 $\text{s}^{-1}\mu\text{M}^{-1}$) varied with MM concentration and, therefore, likely corresponds to binding. The other (20 $\text{s}^{-1}$) was independent of MM concentration and may be related to nucleotide pocket rearrangement due to domain opening. The error reflects the standard deviation of the coefficient (or intercept) of the linear fit. B) The GTPase reaction was run beyond the linear regime for various initial conditions. C and D) The global fit (solid lines) over all measured points for two models: the hydrolysis rate for heterogeneous MM dimers, i.e. bound to one GDP and one GTP, is set to zero (C), or allowed to be non-zero (D). Two important results are obtained from this analysis: (1) an independent measurement of 6.6 $\text{s}^{-1}\mu\text{M}^{-1}$ for the $\text{k}_{\text{on}}$ of GTP binding to MM, (2) evidence that hydrolysis proceeds equally fast for MM dimers containing either 2 GTP or 1 GTP and 1 GDP. See Section S1.1.2 for further discussion.

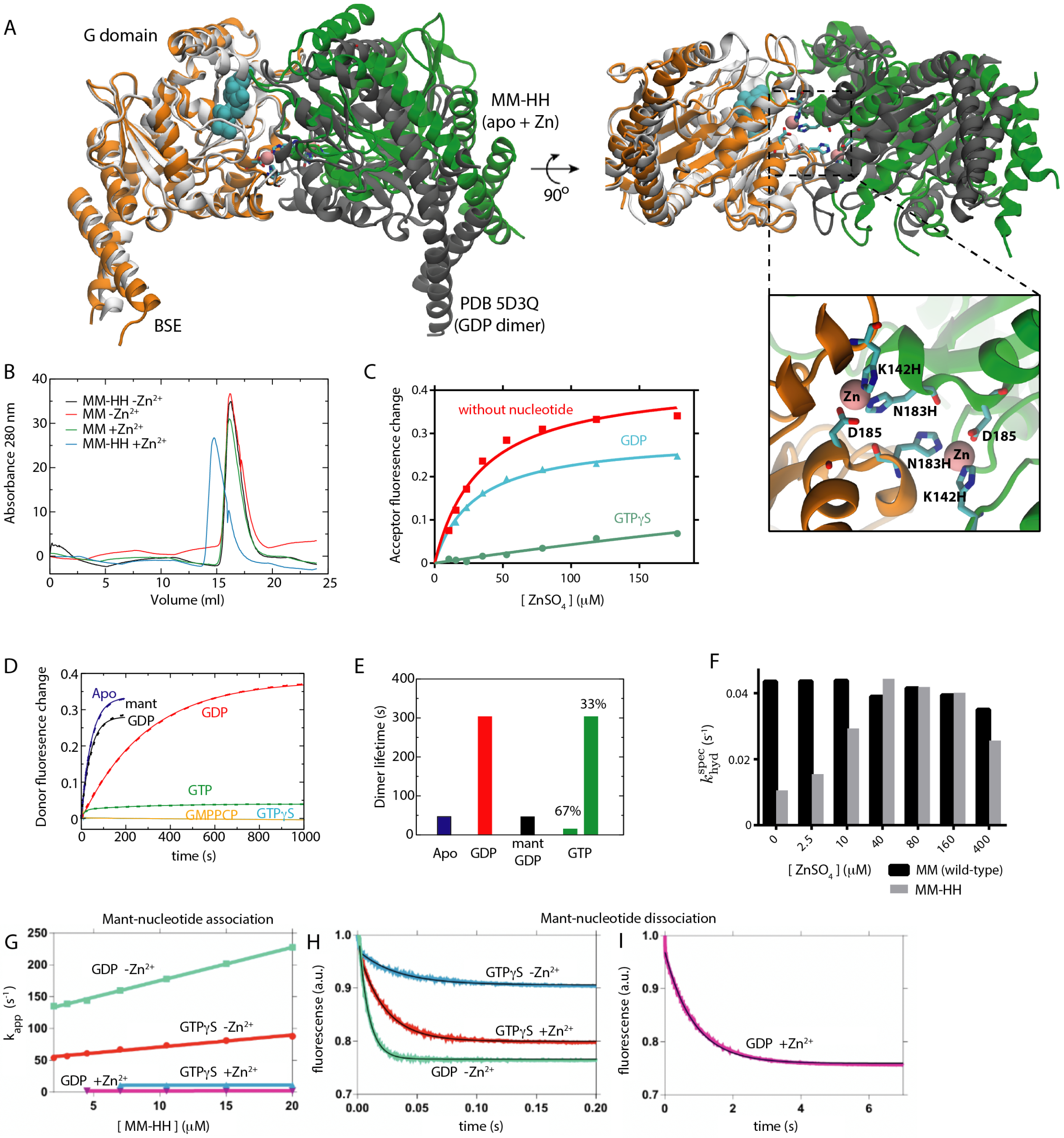

J

| | mantGTPγS -Zn | mantGDP -Zn | mantGTPγS +Zn | mantGDP +Zn |
|---|---|---|---|---|
| $k_{on}$ ($s^{-1}\mu M^{-1}$) | 1.9 ±0.1 | 5.2 ± 0.1 | 0.03 ± 0.01 | 0.019 ± 0.001 |
| $k_{off}$ ($s^{-1}$) | 47 ± 1 | 122 ± 1 | 27 ± 3 | 1.19 ± 0.01 |

Figure S2: Double histidine mutant MM-HH (K142H, N183H) forms a $Zn^{2+}$-stabilized G-interface. A) We confirmed the correct G interface orientation of GDP-bound MM-HH dimers by crystal structure analysis (see also Table S2). Two views on the crystal structure (orange and green, $Zn^{2+}$ as pink spheres) containing the K142H/N183H mutation compared to a GDP-bound wt structure (white and black, PDB 5D3Q [7]). One GDP molecule is shown as cyan spheres. The orange monomer is fitted to the white monomer, showing a nearly unchanged monomer structure (3.1 Å $C_\alpha$ RMSD). The shift of green versus black reveals a subtle repositioning of the G-interface. Zoom shows the residues coordinating the two $Zn^{2+}$ ions stabilizing the dimer. His142 and His183 from one monomer coordinate $Zn^{2+}$ with Asp185 from the other monomer. $Zn^{2+}$ sites were confirmed by their anomalous signal. Note that the crystal structure contains a dimer in the asymmetric unit, i.e. the dimer is not generated by a crystallographic 2-fold axis. B) Size exclusion chromatography verifying that $Zn^{2+}$ stabilizes a dimer in the MM-HH but not MM. Single histidine mutants did not show a significant shift (data not shown). We thus proceeded with the double mutant. C) Ensemble FRET assay to verify MM-HH association in solution. An additional Cys (T165C) was introduced in MM-HH to provide an attachment site for FRET dyes Alexa Fluor 488 or 594. Acceptor fluorescence was plotted over $ZnSO_4$ concentration. In the absence of nucleotide and presence of 0.5 mM GDP, $Zn^{2+}$-dependent assembly was observed whereas 0.5 mM GTP$\gamma$S greatly reduced assembly. GTP and its analogs consistently interfered with the formation of the $Zn^{2+}$ bridge (Figure S2C-E), which precluded further analysis of the dimer association kinetics. D,E) Stopped-flow data to measure the dissocation of HH-MM dimers (D) and resulting rate constants (E). Labelled MM-HH dimers were incubated in the presence of 0.5 mM of the indicated nucleotides in the presence of 160 μM $Zn^{2+}$ and then mixed with an excess of unlabeled MM-HH. Dissociation of MM-HH dimers was determined by monitoring the increase of donor fluorescence. GDP stabilized the dimer resulting in a 6-fold increase in life-time compared to the apo form, whereas mant-GDP did not (see also panel I and J). No dissociation was observed for GTP analogs because these nucleotides mostly precluded dimerization. For GTP, a small dissociation signal was observed that had two components. A fast dissociation rate likely corresponds to the dissociation of residual GTP-bound dimers, whereas the second rate matched GDP-bound dimer dissociation and may thus represent a long-lived post hydrolysis species. F) Specific hydrolysis rate plotted as a function of $Zn^{2+}$ concentration. MM-HH shows GTPase activity, although reduced by a factor of 4 in the absence of $Zn^{2+}$. Addition of $Zn^{2+}$ enhances GTPase activity, presumably by promoting dimerization, although never above the wt level. This is further evidence that dimerization of the MM-HH construct is destabilized by GTP. G,H,I) Stopped flow measurements to determine the on and off rates of mant-nucleotide binding to MM-HH in either the presence (+$Zn^{2+}$) or absence (-$Zn^{2+}$) of $Zn^{2+}$. On rates were determined by varying the MM-HH concentration with a constant concentration of mant-nucleotide (G). Panels H and I show off-rate determination by incubating MM-HH with nucleotide and then mixing with unlabeled nucleotide. J) Table with the fitted rate constants. Note that the mant-GDP dissociation rate is greatly reduced upon dimerization (e.g. in the presence of $Zn^{2+}$), whereas in the absence of $Zn^{2+}$(e.g. in the monomeric state), the rates are comparable to the wt (Figure 1D).

A

| Crystal structure | | | smFRET | | Simulation | |
|---|---|---|---|---|---|---|
| Conformation | Å | FRET | Conformation | FRET | Conformation | FRET |
| open (GMPPCP) | 85 | 0.06 | open (GTP) | 0.16 | open (GMPPCP) | 0.19 |
| closed (GDP+AlF$_4$) | 59 | 0.37 | mid (GTP/GTPγS) | 0.36 | | |
| closed (apo) | 43 | 0.79 | closed (apo) | 0.77 | closed (apo) | 0.79 |

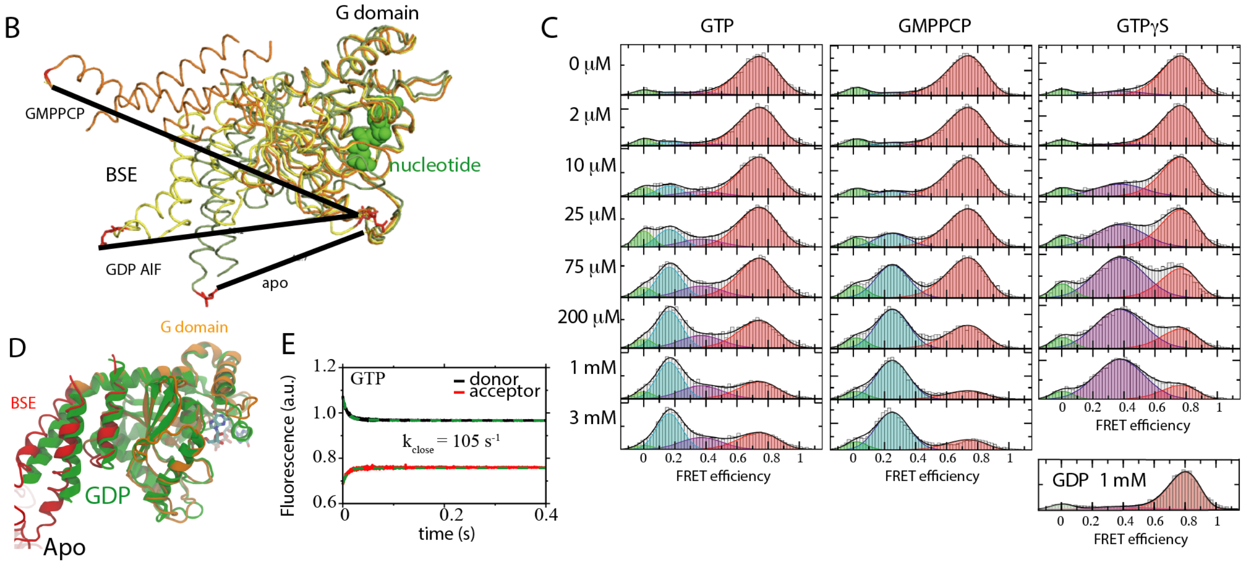

Figure S3: Comparison between smFRET efficiency and the molecular structure of the MM. A) Summary of distance-type comparisons. Distances $d$ between the labeling sites T165C and R318C are measured in the following crystal structures 3ZYC (GMPPCP), 2X2E (GDP·AlF$_4^-$), and 3SNH (apo). The corresponding FRET efficiency was computed using $E = (1 + (d/d_0)^6)^{-1}$ and $d_0 = 5.4$ nm [8,9]. The smFRET entries are taken from the experimental peaks (Figure 4A). The simulation entries are computed by averaging $E$ over the simulation trajectory, where $d$ is defined by the distance between FRET probes. B) Overlay of three crystal structures from panel A. Lines show the displacement of FRET dye labeling sites. C) Full titrations for the nucleotides that induce an opening transition. GDP reproduced for clarity. D) Overlay of the apo (PDB 3SNH) and GDP-bound (PDB 5D3Q) MM crystal structures shows that the BSE is slightly closer to the G domain in the GDP-bound form. This is in agreement with the smFRET histograms. E) Ensemble kinetics of MM closing (see Figure 2D for opening) measured by mixing GTP-bound doubly-labeled MM with buffer containing no GTP. Note that the measured closing rate encompasses two processes: domain closure and GTP dissociation.

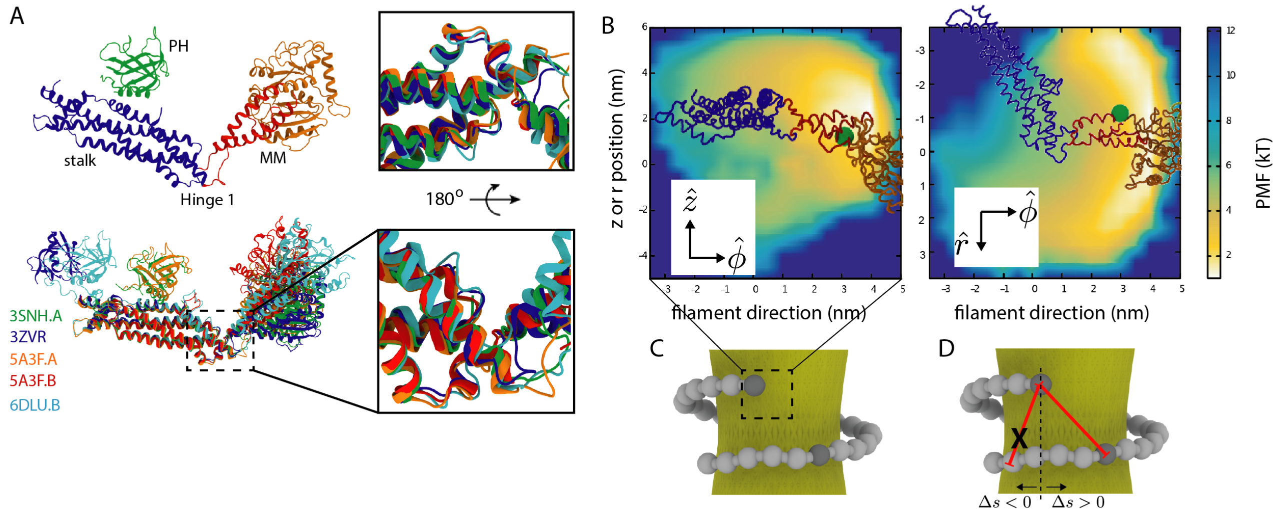

Figure S4: Hinge 1 biases MM orientation. A) Dynamin monomer with the two loop regions comprising hinge 1 is shown. Hinge 1 connects the MM to the stalk. Overlay of five atomic resolution structures [6, 10–12] (PDB codes are shown in the figure) of the dynamin monomer aligned by the stalk domain. The hinge 1 structure and the relative MM/stalk configuration are consistent between the structures implying that this is the preferred configuration of hinge 1. B) A structure-based MD simulation of an upper rung MM attached to its stalk shows the asymmetry of the MM orientation under thermal fluctuations. Heat map plots the potential of mean force (PMF) for the position of Pro294 in hinge 2 (green sphere) relative to the stalk domain, which is frozen during the simulation. For plotting, the stalk is placed at the position of an upper rung stalk in the non-constricted state (Figure 4A). Two different views of the distribution in the $r$-plane or the $z$-plane. Clearly, hinge 1 forces the orientation of the MM (monitored by position of Pro294) to preferentially point to the right (i.e. $\Delta s > 0$ in Figure S4). C) Geometry of the simulated heat map in the context of the filament. D) Hinge 1 positioning, i.e. toward the right in the upper rung or to the left in the lower rung, has the effect of preventing the initial formation of MM dimers with $\Delta s < 0$. This is important because such dimers would have produced the expanding torque.

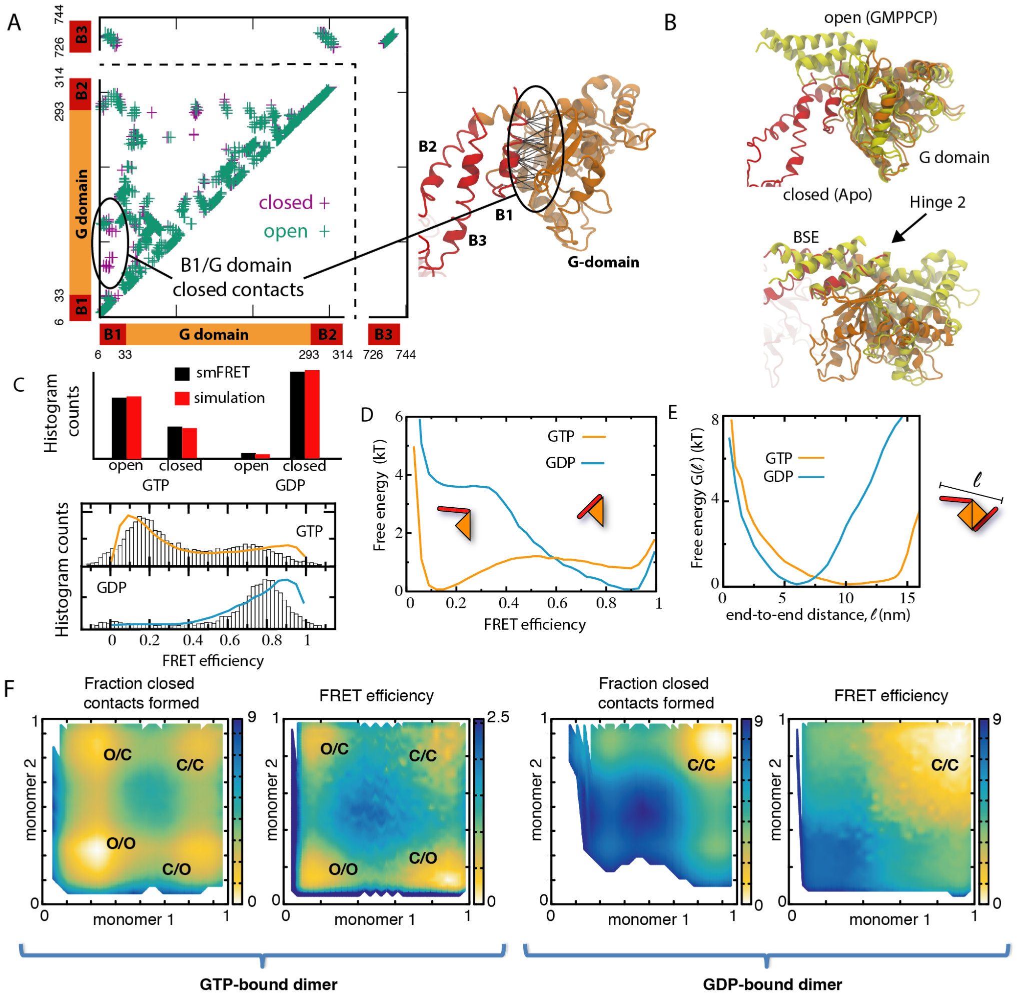

Figure S5: Input to and results from the dual-basin structure-based model. Additional explanatory text is in Section S1.2. A) Native contact map comparison between the apo (closed) structure and the GMPPCP (open) structure. The contact map indicates residue pairs that have atoms in close proximity (within 6 Å [13]) in the crystal structures. The contact maps are essentially identical except for the additional G domain/BSE interface in the closed structure. The dual-basin model weighting factor $\lambda$ allows the stability of these unique closed contacts to be varied in order to tune the relative stability between the open and closed states. B1 – helix 1, B2 – helix 2, B3 – helix 3 in BSE. B) The open GMPPCP structure fitted to the closed apo structure via the G domain (upper) or the BSE (lower). In both cases the domain architecture is very similar, the conformational change is limited to the shifts within hinge 2. C) (top) The weighting factor in the dual-basin simulations is chosen such that the simulated FRET histograms agree with the experimental smFRET distributions on a coarse level. The open (closed) state was defined as having FRET $< 0.5$ ($> 0.5$). (bottom) Complete experimental and simulated FRET histograms are compared; note that a detailed quantitative agreement is not expected here due to averaging on the experimental level and to simplifications within the simulation. D) Free energy as a function of the simulated FRET efficiency for MM monomers. E) Connecting two monomers by the G interface allows us to determine the dimer free energy as a function of its reaction coordinate $\ell$ (the end-to-end distance) in two nucleotide states. F) Two-dimensional free energy landscapes from MM dimer simulations plotted with different monomer-based reaction coordinates: (left) the fractions of unique closed contacts or (right) the simulated FRET efficiencies for the two monomers. The energy units are $k_B T$; the MM dimer states are denoted as, e.g., O/C (open/closed). The unique closed contacts are defined by the circled group in Panel A. A contact was considered as formed if the distance between the corresponding residue pair was less than 1.2 times its distance in the closed (GDP-bound) crystal structure. Computing the simulated FRET efficiency is described in the Methods.

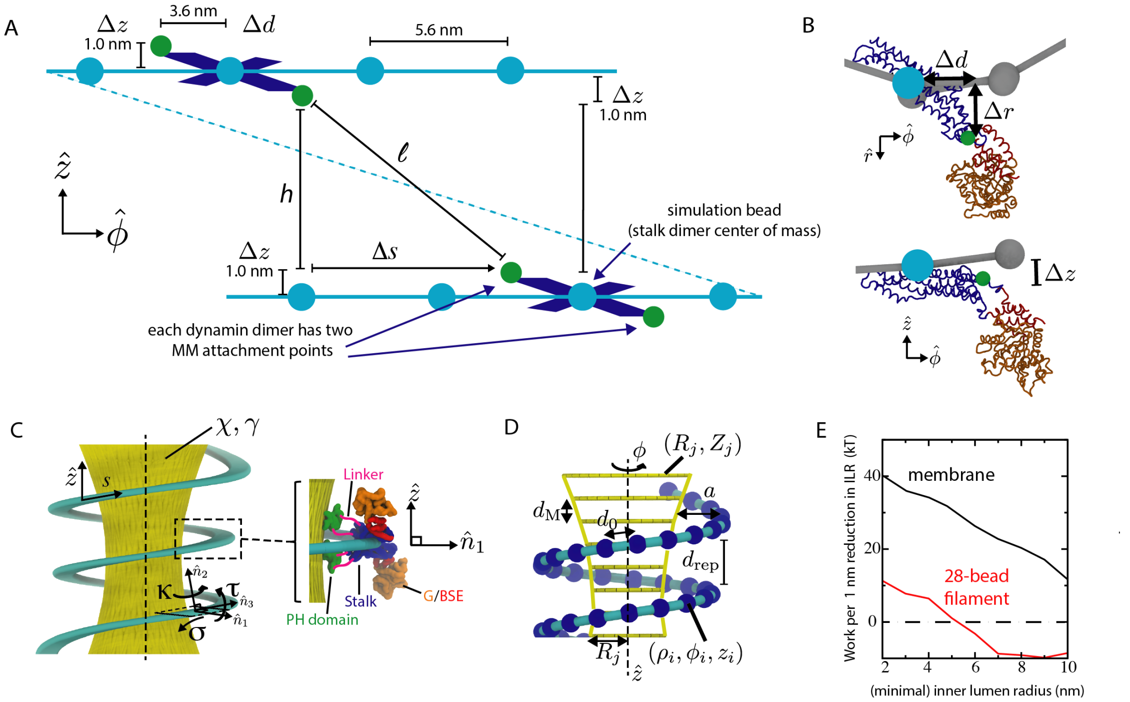

Figure S6: Simulation geometry and definition of variables in the elastic filament model. A) Geometry of the force application. The position of a simulation bead (cyan) represents the center of mass of a dynamin stalk dimer. The position of the MM attachment point (hinge 1) can be determined relative to the dimer center of mass from the crystal structure of the dynamin dimer. Thus, each simulation bead, centered at $(r_0,\phi_0,z_0)$ has two virtual sites (green) associated with it, one at $(r_0+\Delta r,\phi_0+\Delta d/(r_0+\Delta r),z_0-\Delta z)$ and the other at $(r_0+\Delta r,\phi_0-\Delta d/(r_0+\Delta r),z_0+\Delta z)$, with $\Delta r = 3.7$ nm, $\Delta d = 3.6$ nm, and $\Delta z = 1.0$ nm. A force applied at hinge 1 is treated as a force applied at its center of mass. Note that even though the simulation snapshots depict the cross-dimer attachment points as being the centers of the beads, in the simulation the distance $\ell$ is actually calculated between the MM attachment points (green beads, hinge 1). B) Orientation of a dynamin monomer within the filament showing the displacements between the center of mass (cyan dot) and hinge 1 (green dot). These displacements are held constant during the simulation, which keeps the stalk dimer rigid. C) The structural part of the dynamin helix is formed by the stalk filament, which is treated as an oriented elastic ribbon with curvature $\kappa$, twist $\tau$, and geodesic curvature $\sigma$. The membrane tube is an axisymmetric elastic sheet with stiffness $\chi$ and under tension $\gamma$. This underlying filament+membrane model is summarized in the Methods or see [14] for a full discussion. D) For numerical simulations, both the filament and membrane are discretized. The filament has a natural discretization of a single bead per dynamin dimer (blue bead) and, thus, the beads are spaced at $d_0 = 5.6$ nm intervals. The membrane tube is modeled as a stack of cylinders (yellow rungs), each with an axial width of $d_{\mathrm{M}} = 4$ nm. E) Comparison of the effective stiffness, i.e. the amount of work needed to reduce the ILR by 1 nm, between a membrane tube with $\chi = 24\ \mathrm{k_BT}$ and its constricting dynamin stalk filament. The total energy of the elastic system is divided between the filament and the membrane. Until the ILR is below 5 nm, the filament contributes to constriction. Below 5 nm, the motors must work against both the filament and the membrane, and at the tightest ILR, deforming the filament requires a significant portion of the total work.

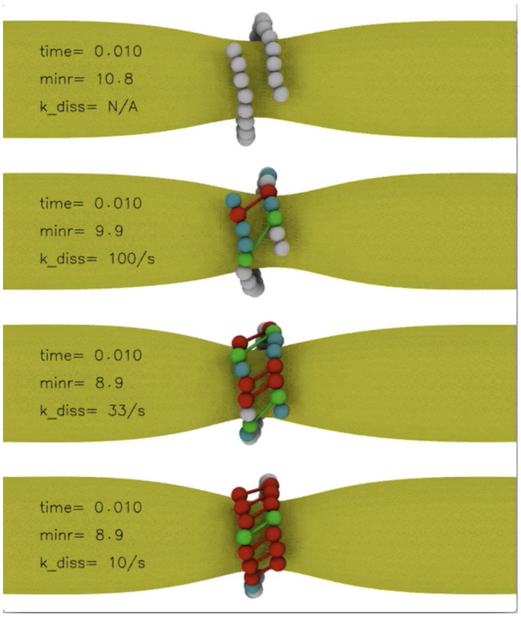

Video S1: Dependence of constriction dynamics on MM dimer dissociation rate $k_{\rm diss}$. Decreasing $k_{\rm diss}$ increases the duty ratio $\omega$ and, therefore, the fraction of bridging cross-dimers that are simultaneously present. A larger set of cross-dimers generates a stronger torque, but their more slow dissociation leads to development of a larger number of blocking links that decrease the constriction speed. The filament contains $N = 28$ dynamin dimers. The top panel shows the dynamics in the absence of GTP. The other three panels show the first 2 seconds of simulations after the addition of GTP for $k_{\rm diss} = (100, 33, 10)$ s$^{-1}$. White beads (GTP-bound monomers or apo in the top panel), cyan beads (GDP-bound monomers), green beads and links (GTP-bound dimers), red beads and links (GDP-bound dimers). Recall that each bead is a dynamin dimer and has two associated MMs. The color of the bead corresponds to only one of the associated MMs, the coloring for the right rung indicates the state of the MMs pointing to the left and for the left rung indicates the state of the MMs pointing to the right.

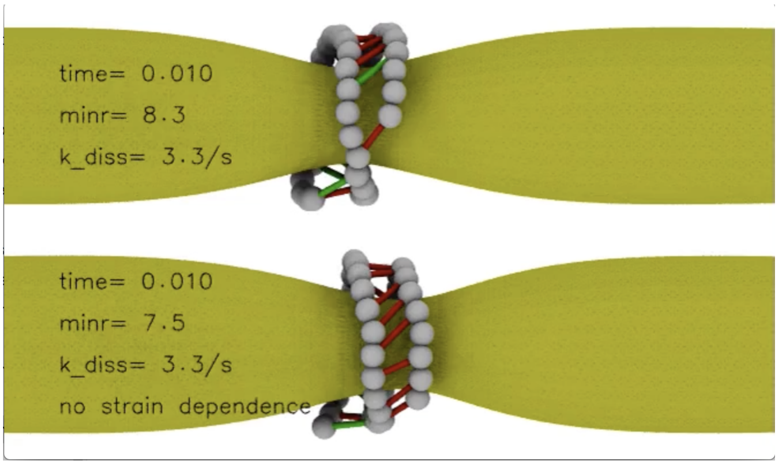

Video S2: Strain-dependent dissociation of cross-dimers reduces blocking and accelerates constriction. Upper panel, strain dependence; lower panel, no strain dependence. The first 2 seconds after the addition of GTP are shown, $N = 40$. Green/red links correspond to GTP/GDP-bound cross-dimers. Bead coloring is not indicative of the state.

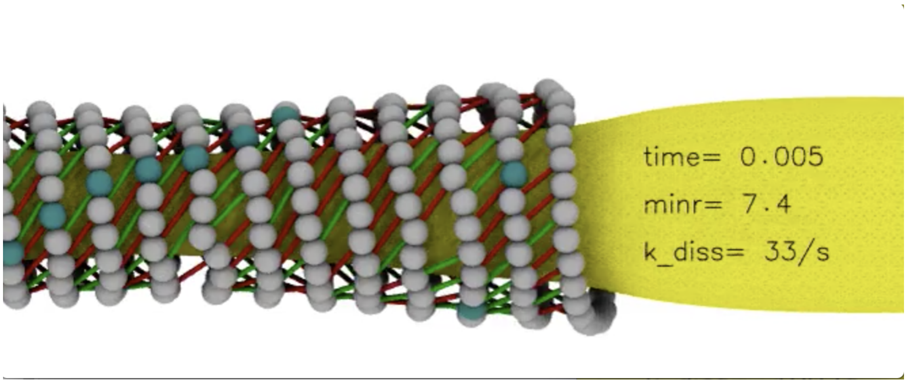

Video S3: Constriction process in a long filament ($N = 900$) under strain dependence of the dimer dissociation rate. Only $\sim 25\%$ of the filament is displayed. The first 600 ms of a simulation are shown. Every 20th bead is colored cyan as a guide to eye. Green/red links correspond to GTP/GDP-bound dimers. Bead coloring is not indicative of the state.

| **Data Collection** | |
|---|---|
| Space group | P 1 $2_1$ 1 |
| Cell dimensions | |
| *a, b, c* (Å) | 70.69 69.55 74.28 |
| α, β, γ (°) | 90 113.15 90 |
| Wavelength | 1.2825 Å |
| Resolution (Å)* | 47-2.0 (2.071 – 2.0) |
| No. reflections | 44,543 (4,424) |
| $I/\sigma I$* | 15.73 (1.24) |
| Completeness (%)* | 99.8 (100) |
| $R_{meas}$ (%) | 5.4 (76.2) |
| $CC_{1/2}$ | 0.998 (0.71) |
| **Refinement** | |
| $R_{work}$ / $R_{free}^{\S}$ (%) | 18.6/22.3 (29.4/30.5) |
| No. atoms | |
| Protein | 4,997 |
| Ligands | 15 |
| Water | 0 |
| B-factors | |
| Protein | 73.0 |
| Ligands | 89.8 |
| R.m.s deviations | |
| Bond lengths (Å) | 0.014 |
| Bond angles (°) | 1.26 |
| Ramachandran | |
| Favored (%) | 96 |
| Allowed (%) | 3.7 |
| Outliers (%) | 0.32 |
| Clash score | 6.8 |
| No. TLS groups | 40 |

* Statistics for the highest-resolution shell are shown in parentheses

& 5% of reflections were excluded from refinement

Table S2: Crystallographic data of the nucleotide-free MM-HH (K142H, N183H) structure.

### S2 Supplementary Methods

#### S2.1 Analytical SEC

To characterize $Zn^{2+}$-mediated dimerization of single- or double-histidine mutants of the MM, SEC was performed on a Superdex S200 column 10/300 GL equilibrated with buffer D (20 mM HEPES-NaOH, pH 7.5, 150 mM NaCl, 5 mM KCl, 0.2 mM $ZnSO_4$). For complex formation, 10-15 µl of the corresponding protein variant was mixed with buffer D. Following incubation on ice for 10 min, 100 µl of a 4 mg/ml solution was applied to SEC.

#### S2.2 Protein crystallization and structure determination

For crystallization, MM-HH (MM with mutations K142H, N183H) dimer at 12 mg/ml was mixed in a 1:1 ratio with reservoir solution containing 0.1 M Tris-HCl, 0.25 M KBr, 30% (w/v) PEG MME 2000. Crystals were grown by sitting drop vapor diffusion at 20 °C. A dataset was collected from a single cryo-cooled crystal at BESSY synchrotron beamline BL14.1 (Berlin, Germany). Diffraction data to a maximal resolution of 2.0 Å were included for processing by the XDS software package [15]. The structure was solved by molecular replacement using the coordinates for GTPase-BSE of PDB entry 5D3Q using Phaser [16] within the software package Phenix [17]. Refinement and model validation were performed with Phenix. Model building was performed manually with COOT [18]. Six zinc ions were located based on the coordination spheres and positive difference densities after placement of water molecules at these positions. The model was deposited in the PDB database under accession number 6S9A.

#### S2.3 $Zn^{2+}$-dependent MM dimerization

For determining the concentration of $Zn^{2+}$ required to promote dimerization, MM-HH(K142H, N183H) including T165C was separately labeled via Cys165 with either Alexa Fluor 488 or 594 dyes. The labelled proteins were mixed in a 1:2 molar ratio (Alexa Fluor 488: Alexa Fluor 594) in 20 mM HEPES-NaOH, pH 7.5, 150 mM KCl, 4 mM $MgCl_2$. In order to measure Kd values, 5 µM of the labelled protein mixture was incubated with different concentrations of $ZnSO_4$ (177.8, 118.5, 79.0, 52.7, 35.1, 23.4 and 15.6 µM) and incubated for 10 min on ice. The resulting FRET signal was recorded on a Teacan1000pro plate reader ($\lambda_{ex}$= 465 nm and $\lambda_{em}$=616 nm).

To account for $ZnSO_4$-mediating quenching effects on the AF488 dye, a correction factor was calculated: the fluorescence intensity of AF488-labeled protein in the absence of $ZnSO_4$ was divided by the fluorescence value in the presence of each particular $ZnSO_4$ concentration. Subsequently, the background FRET signal in the protein sample without $ZnSO_4$ was subtracted from each measured and corrected data point. FRET data were plotted against $ZnSO_4$ concentration. The titration indicated that dimerization saturated near 200 µM $Zn^{2+}$.

In order to evaluate the effect of nucleotide on dimerization, the same type of experiments were performed in the presence of 160 µM $Zn^{2+}$ and 0.5 mM GTP$\gamma$S or GDP. To exclude an influence of $MgCl_2$ on dimerization, additional experiments were performed in the absence or presence of 2 mM $MgCl_2$.

#### S2.4 Dual-basin energy potential for motor domain MD simulations

The dual-basin SBM was constructed in such a way that both the known open and closed MM structures were explicit energetic minima of it. The open structure was defined by chain A of the GMPPCP-bound MM ($S_O$) [19] and the closed structure by chain A of the GTP-bound MM ($S_C$) [7]. The G domains between the two structures are nearly identical. The structural differences mainly reside in the positioning of the first BSE helix relative to the G domain (Figure S5). The native configuration for the loop connecting the second and third BSE helices (which was unresolved in the crystal structures) was taken as the hinge 1 configuration in the tetramer crystal structure with a 3.8 Å bond inserted between Pro322 and Glu716.

We followed the general approach [20] in defining the dual-basin potential, except for two modifications: (i) a Gaussian-shaped native contact interaction potential that could have two minima was used [21] and (ii) angle and dihedral interactions, differing between the structures by more than a cutoff, were removed. Namely, the native contact maps [13], i.e. those residues that are close in the native structure, were divided into three contact maps (M): (1) the contacts ($M^U_O$) unique for the open state $S_O$ or (2) those ($M^U_C$) unique for the closed state $S_C$ , and (3) the contacts ($M^S$) shared by both structures. All native contacts were given a stabilizing interaction centered at the native distance, the shared contacts had a minimum at both distances. Bond angles between three sequential residues and dihedral angles were considered frustrated and not given stabilizing interactions if the angles differed by more than 20° or 30°, respectively. The remaining angles were biased toward the average angle between the $S_O$ and $S_C$.

The energy potential was

$$V_{\text{dual}}(\lambda) = \sum_{ij\in\text{bonds}} \epsilon_b(r_{ij} - r_{ij}^C)^2 + \sum_{ijk\in\text{angles}} \epsilon_\theta \left(\theta_{ijk} - \frac{\theta_{ijk}^C + \theta_{ijk}^O}{2}\right)^2 + \sum_{ijkl\in\text{dihedrals}} \epsilon_\phi F_D \left(\phi_{ijkl} - \frac{\phi_{ijkl}^C + \phi_{ijkl}^O}{2}\right)$$
$$\sum_{ij\in M_C^U} N_{ij}(\lambda\epsilon_N, r_{ij}^C) + \sum_{ij\in M_O^U} N_{ij}(\epsilon_N, r_{ij}^O) + \sum_{ij\in M^S} N_{ij}(\epsilon_N, r_{ij}^C, r_{ij}^O) + \sum_{ij\notin contacts} \epsilon R_{ij}$$

where the factor $\lambda$ allowed the weight of closed states versus open states to be modulated. Superscript O and C denote that the value is taken from either the open or closed crystal structure. $F_D$ is a typical dihedral-like potential

$$F_D(x) = (1 - \cos x) + 0.5(1 - \cos 3x)$$

and $N_{ij}$ is a native contact interaction consisting of a Gaussian well $G_{ij}$ coupled a $r^{-12}$ repulsion $R_{ij}$ in such a way that fixes the minimum at $(r_0, \epsilon)$ [22].

$$N_{ij}(\epsilon, r_0) = \epsilon[(1 + (1/\epsilon)R_{ij})(1 + G_{ij}(r_0)) - 1]$$

with

$$G_{ij}(r_0) = -e^{(r_{ij}-r_0)^2/2\sigma^2}$$

and

$$R_{ij} = (a/r_{ij})^{12},$$

where $a = 4$ Å sets the bead excluded volume. The dual minimum contact interaction was similarly constructed,

$$N_{ij}(\epsilon, r_0^1, r_0^2) = \epsilon[(1 + (1/\epsilon)R_{ij})((1 + G_{ij}(r_0^1))(1 + G_{ij}(r_0^2))) - 1]$$

The values of the energetic parameters are $\epsilon_b = 20000\epsilon$, $\epsilon_\theta = 40\epsilon$, $\epsilon_\phi = \epsilon$, $\epsilon_N = \epsilon$, where $\epsilon$ is the reduced energy unit.

Simulations were performed using GROMACS 4.5.3 [23] containing code edits implementing Gaussian contact interactions (available at http://smog-server.org). Single basin simulation topologies were generated using the SMOG2 software [24] with the forcefield "SBM_CAgauss" and combined using an in-house script. The temperature (T=0.92 in reduced units) was chosen such that the average $C_\alpha$ atom RMSD in the closed MM agreed between a 100 ns explicit solvent simulation using AMBER99SB-ILDN at 310K and the SBM. The temperature was maintained with stochastic dynamics with coupling constant 0.1 and the time step was 0.0005.

To ensure the sampling of rare states, particularly important for extended conformations of the GDP-bound dimer, umbrella sampling was used. The umbrella coordinate was the number of unique closed native contacts made between the BSE and G domain and, thus, ensured sampling from fully open to fully closed. The dimer simulations had two independent umbrella coordinates, one for each monomer. Unbiased histograms were obtained using the weighted histogram analysis method [25], and were used to calculate free energies.

#### S2.5 Elastic description of the filament and membrane tube

The code used to implement the constriction model (written in Java) is publicly available at https://bitbucket.org/jknoel/constrictionsimulation. A detailed formulation and analysis of the model for a dynamin filament on a membrane tube, in absence of the motor activity, was given in the previous publication [14]. The dynamin filament was represented as an elastic polymer with each bead corresponding to a dynamin dimer. Stiffness constants for various elastic deformations of the polymer (i.e., the stretch $k$, the normal curvature $\kappa$, the twist $\tau$, and the geodesic curvature $\beta$) were obtained from molecular simulations of dynamin tetramers or short filaments. The geometries of the continuous elastic system and its discretization are shown in Figure S6C,D. The elastic energy of the filament $E_F$ was

$$E_F = d_0 \sum_{i=1}^{N} \left[ \beta \left( \frac{\kappa_i}{\kappa_i^2 + \tau_i^2} \frac{\partial \tau_i}{\partial s} - \frac{\tau_i}{\kappa_i^2 + \tau_i^2} \frac{\partial \kappa_i}{\partial s} \right)^2 + \alpha_\kappa (\kappa_i - \kappa_0)^2 \right.$$
$$\left. + \alpha_\tau (\tau_i - \tau_0)^2 + k(|\vec{r}_i - \vec{r}_{i-1}| - d_0)^2 + E^{\text{repulsion}} \right] \quad \text{(S9)}$$

Note that $E^{\text{repulsion}}$ prevented the local pitch, i.e. axial distance between nearest neighbor filaments, from going below 8 nm.

The membrane tube description was based on an axially symmetric continuous Helfrich elastic membrane with stiffness $\chi$ and under tension $\gamma$. It was discretized into 4 nm disks for numerical simulations. The elastic energy of the membrane $E_{\mathrm{M}}$ was

$$E_{\mathrm{M}} = \pi d_{\mathrm{M}} \left[ \sum_{j=1}^{M} \left( \frac{\chi}{R_j} + 2R_j\gamma \right) + \chi \sum_{j=3}^{M-2} R_j \left( \frac{d^2[R_j]}{dz^2} \right)^2 \right] \tag{S10}$$

The coupling energy between the filament and the membrane was

$$E_{\mathrm{int}} = \frac{d_{\mathrm{M}}}{2} \epsilon_{\mathrm{int}} \sum_{i=1}^{N} \left[ r_i - a - \bar{R}(i) \right]^2 \tag{S11}$$

The separation between the center of the filament and the center of the membrane bilayer was $a = 8.5$ nm, which was taken from cryo-EM structures. The inner lumen radius (ILR) was defined as the membrane radius minus 2 nm (radius of dynamin stalk minus 10.5 nm), which assumed a 4 nm wide bilayer. The time evolution equations (given in publication [14]) locally conserved the membrane area and the time scale was defined by a single lipid diffusion coefficient of 1 $\mathrm{nm}^2/\mu\mathrm{s}$. The filament beads were in contact with a thermal bath at 310 K, while thermal fluctuations in the membrane came only through contact with the beads. The motion of the beads was described by Langevin equations, see [14]. The mobility of a filament bead bound to the membrane, 1 $\mathrm{nm}^2/\mu\mathrm{s}$, was chosen to approximate the diffusion coefficient of single lipids within a bilayer. The lipid flows within the membrane were controlled by the same time scale [14]. The time step of 10 ns in numerical simulations was limited by the membrane description.

### References

[1] Chappie, J. S, Acharya, S, Leonard, M, Schmid, S. L, & Dyda, F. (2010) G domain dimerization controls dynamin's assembly-stimulated GTPase activity. *Nature* **465**, 435–440.

[2] Noel, J. K. (2012) Development and Application of All-Atom Structure-based Models for Studying the Role of Geometry in Biomolecular Folding and Function. *UCSD* pp. 1–153.

[3] Giri Rao, V. V. H & Gosavi, S. (2016) Using the folding landscapes of proteins to understand protein function. *Curr. Opin. Struct. Biol.* **36**, 67–74.

[4] Sun, L, Noel, J. K, Sulkowska, J. I, Levine, H, & Onuchic, J. N. (2014) Connecting Thermal and Mechanical Protein (Un)folding Landscapes. *Biophys. J.* **107**, 2941–2952.

[5] Schlosshauer, M & Baker, D. (2004) Realistic protein-protein association rates from a simple diffusional model neglecting long-range interactions, free energy barriers, and landscape ruggedness. *Protein science : a publication of the Protein Society* **13**, 1660–1669.

[6] Faelber, K, Posor, Y, Gao, S, Held, M, Roske, Y, Schulze, D, Haucke, V, Noé, F, & Daumke, O. (2011) Crystal structure of nucleotide-free dynamin. *Nature* **477**, 556–560.

[7] Anand, R, Eschenburg, S, & Reubold, T. F. (2016) Crystal structure of the GTPase domain and the bundle signalling element of dynamin in the GDP state. *Biochemical and Biophysical Research Communications* **469**, 76–80.

[8] Schuler, B, Lipman, E. A, & Eaton, W. A. (2002) Probing the free-energy surface for protein folding with single-molecule fluorescence spectroscopy. *Nature* **419**, 743–747.

[9] Schuler, B & Hofmann, H. (2013) Single-molecule spectroscopy of protein folding dynamics–expanding scope and timescales. *Curr. Opin. Struct. Biol.* **23**, 36–47.

[10] Ford, M. G. J, Jenni, S, & Nunnari, J. (2011) The crystal structure of dynamin. *Nature* **477**, 561–566.

[11] Reubold, T. F, Faelber, K, Plattner, N, Posor, Y, Ketel, K, Curth, U, Schlegel, J, Anand, R, Manstein, D. J, Noé, F, Haucke, V, Daumke, O, & Eschenburg, S. (2015) Crystal structure of the dynamin tetramer. *Nature* **525**, 404–408.

[12] Kong, L, Sochacki, K. A, Wang, H, Fang, S, Canagarajah, B, Kehr, A. D, Rice, W. J, Strub, M.-P, Taraska, J. W, & Hinshaw, J. E. (2018) Cryo-EM of the dynamin polymer assembled on lipid membrane. *Nature* pp. 1–17.

[13] Noel, J. K, Whitford, P. C, & Onuchic, J. N. (2012) The Shadow Map: A General Contact Definition for Capturing the Dynamics of Biomolecular Folding and Function. *J. Phys. Chem. B* **116**, 8692–8702.

[14] Noel, J. K, Noé, F, Daumke, O, & Mikhailov, A. S. (2019) Polymer-like Model to Study the Dynamics of Dynamin Filaments on Deformable Membrane Tubes. *Biophys. J.* **117**, 1870–1891.

[15] Kabsch, W. (2010) Integration, scaling, space-group assignment and post-refinement. *Acta Crystallogr., Sect. D: Biol. Crystallogr.* **66**, 133–144.

[16] McCoy, A. J, Grosse-Kunstleve, R. W, Adams, P. D, Winn, M. D, Storoni, L. C, & Read, R. J. (2007) Phaser crystallographic software. *J. Appl. Crystallogr.* **40**, 658–674.

[17] Adams, P. D, Afonine, P. V, Bunkóczi, G, Chen, V. B, Davis, I. W, Echols, N, Headd, J. J, Hung, L.-W, Kapral, G. J, Grosse-Kunstleve, R. W, McCoy, A. J, Moriarty, N. W, Oeffner, R, Read, R. J, Richardson, D. C, Richardson, J. S, Terwilliger, T. C, & Zwart, P. H. (2010) PHENIX: a comprehensive Python-based system for macromolecular structure solution. *Acta Crystallogr., Sect. D: Biol. Crystallogr.* **66**, 213–221.

[18] Emsley, P & Cowtan, K. (2004) Coot: model-building tools for molecular graphics. *Acta Crystallogr., Sect. D: Biol. Crystallogr.* **60**, 2126–2132.

[19] Chappie, J. S, Mears, J. A, Fang, S, Leonard, M, Schmid, S. L, Milligan, R. A, Hinshaw, J. E, & Dyda, F. (2011) A pseudoatomic model of the dynamin polymer identifies a hydrolysis-dependent powerstroke. *Cell* **147**, 209–222.

[20] Whitford, P. C, Miyashita, O, Levy, Y, & Onuchic, J. N. (2007) Conformational transitions of adenylate kinase: Switching by cracking. *J. Mol. Biol.* **366**, 1661–1671.

[21] Noel, J. K, Schug, A, Verma, A, Wenzel, W, Garcia, A. E, & Onuchic, J. N. (2012) Mirror images as naturally competing conformations in protein folding. *J. Phys. Chem. B* **116**, 6880–6888.

[22] Lammert, H, Schug, A, & Onuchic, J. N. (2009) Robustness and generalization of structure-based models for protein folding and function. *Proteins: Struct., Funct., Bioinf.* **77**, 881–891.

[23] Pronk, S, Páll, S, Schulz, R, Larsson, P, Bjelkmar, P, Apostolov, R, Shirts, M. R, Smith, J. C, Kasson, P. M, van der Spoel, D, Hess, B, & Lindahl, E. (2013) GROMACS 4.5: a high-throughput and highly parallel open source molecular simulation toolkit. *Bioinformatics* **29**, 845–854.

[24] Noel, J. K, Levi, M, Raghunathan, M, Lammert, H, Hayes, R. L, Onuchic, J. N, & Whitford, P. C. (2016) SMOG 2: A Versatile Software Package for Generating Structure-Based Models. *PLOS Comput. Biol.* **12**, e1004794.

[25] Kumar, S, Rosenberg, J, Bouzida, D, & Swendsen. (1992) The weighted histogram analysis method for free-energy calculations on biomolecules. I. The method. *J. Comput. Chem.* **13**, 1011.
